# Combined inhibition of AIF/CHCHD4 interaction and GLS1 to exploit metabolic vulnerabilities in pediatric osteosarcoma

**DOI:** 10.64898/2026.04.03.716303

**Authors:** Hong-Toan Lai, Tran Ngoc Anh Nguyen, Maria Eugénia Marques da Costa, Romain Fernandes, Daniela Dias-Pedroso, Sylvère Durand, Guido Kroemer, Reynand Jay Canoy, Liuba Mazzanti, Yegor Vassetzky, Nathalie Gaspar, Antonin Marchais, Birgit Geoerger, Tâp Ha-Duong, Catherine Brenner

## Abstract

Osteosarcoma is a malignant bone tumor with a high risk of metastatic relapse and poor outcomes due to primary and acquired chemoresistance. This highlights the medical need to develop effective targeted approaches to overcome chemoresistance. Recent studies have revealed the roles of metabolic reprogramming and mitochondria-nucleus crosstalk in osteosarcoma progression, indicating the potential of these cellular processes as therapeutic targets. The complex formed by mitochondrial apoptosis-inducing factor (AIF) and coiled-coil-helix-coiled-coil-helix domain-containing protein 4 (CHCHD4) orchestrates the import and oxidative folding of cysteine-rich, nuclear-encoded proteins, thereby regulating key mitochondrial functions and metabolism. Here, we identified mitoxantrone as an inhibitor of the AIF/CHCHD4 mitochondrial import machinery and revealed a new mitoxantrone-induced metabolic vulnerability in some osteosarcoma cell line models, characterized by intracellular glutamine accumulation and an increase in nucleotide synthesis. As a result, synergy was found between mitoxantrone and the glutaminase inhibitor telaglenastat in both *in vitro* and *in vivo* osteosarcoma models. Collectively, our findings position the AIF/CHCHD4 complex as a druggable therapeutic target and provide a combination strategy for mitoxantrone/telaglenastat treatment to overcome metabolic adaptations and chemoresistance in osteosarcoma.

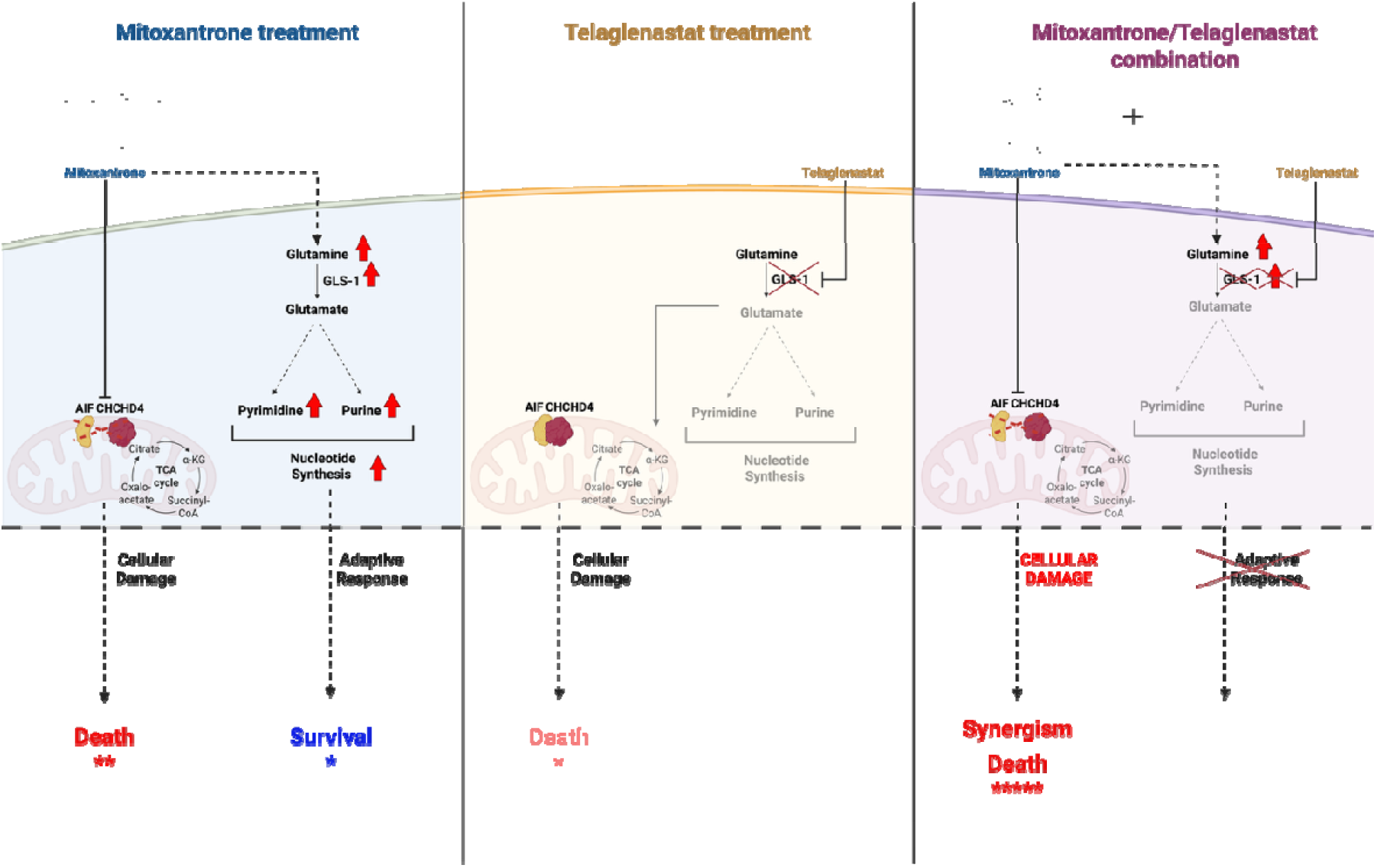

## Introduction

Osteosarcoma is a bone cancer prevalent in adolescents and young adults, with approximately 0.2 to 3 cases per million diagnosed annually in Europe^1^. Patients with osteosarcoma have a high tendency for metastasis, which is commonly observed in the lung or other bones. The exact cause of the disease remains unknown, although certain genetic mutations and hereditary disorders might increase the incidence. Patients diagnosed with localized osteosarcoma have a 5-year overall survival rate of around 75 %; however, for those with metastases, this rate decreases to only 20 % ^2^. First-line standard treatment for osteosarcoma has not substantially changed since the late 1970s, which includes surgical resection of the primary tumor and metastatic localizations, combined with multi-agent chemotherapy ^1 2^. However, the introduction of novel therapies has failed in the last decades and outcomes for patients with metastatic or relapsed osteosarcoma remain dismal, underscoring the urgent need for novel therapeutic approaches. Recent studies highlight metabolic reprogramming as a hallmark of osteosarcoma progression and therapy resistance, positioning tumor metabolism as a potential source of therapeutic vulnerabilities^3 4 5^.

Given the central role of mitochondria in metabolism, thus mediating cancer initiation and progression, this organelle is recognized as plausible targets for antineoplastic drug development^6 7^. We have recently reported a novel approach is to target the mitochondrial apoptosis-inducing factor/coiled-coil-helix-coiled-coil-helix domain containing 4 (AIF/CHCHD4) complex machinery that regulates the assembly of electron transport chain (mETC) and the correct functioning of the mitochondria^8^. More specifically, the CHCHD4-dependent import machinery, also known as the disulfide relay system, is responsible for the folding and oxidation of nucleus-encoded cysteine-rich proteins, facilitating the formation of intramolecular disulfide bonds, and thereby retaining them in the intermembrane space (IMS) to maintain different vital mitochondrial functions ^9^. Thus, pharmacological disruption of AIF/CHCHD4 interaction has recently been demonstrated to inhibit the proliferation of osteosarcoma cell lines and patient-derived xenografts (PDXs) and induce cell death *in vitro*, as well as reduce tumor burden *in vivo*, through rewiring energetic metabolism^8^. Thus, we explored the AIF/CHCHD4 inhibition-induced metabolic changes for the design of novel combinatory cancer treatment approaches.

Here, we identified mitoxantrone, an anthracenedione chemotherapeutic agent, as an AIF/CHCHD4 interaction inhibitor. Initially designed as an analog of doxorubicin, mitoxantrone is a DNA intercalator that provokes DNA strand breaks. It also inhibits topoisomerase II, an enzyme responsible for DNA unwinding and the repair of damaged DNA. mitoxantrone is currently used in chemotherapy regimens for the treatment of breast cancer, non-Hodgkin lymphomas, acute myeloid leukemia, and as a palliative treatment for advanced hormone-resistant prostate cancer ^10^.

To target the metabolic vulnerability of osteosarcoma, we investigated here the interest of repositioning mitoxantrone for osteosarcoma treatment alone and in combination. We observed that (i) mitoxantrone inhibits AIF/CHCHD4 interaction, (ii) mitoxantrone-induced metabolic reprogramming creates a therapeutically exploitable metabolic vulnerability in osteosarcoma and (iii) the combination of mitoxantrone and glutaminase (GLS1) inhibitor telaglenastat synergistically reduces tumor growth in osteosarcoma models *in vitro* and *in vivo*.

## Material and methods

### Chemicals

Telaglenastat (CB-839) (MedChemExpress, #HY-12248) and mitoxantrone (Sigma-Aldrich, #M6545) were purchased from commercial suppliers. Both mitoxantrone and telaglenastat were dissolved in dimethylsulfoxid (DMSO) at 10 mM stock solutions and stored at-20°C for *in vitro* experiments. For *in vivo* experiments, mitoxantrone was dissolved in phosphate-buffered saline (PBS) and telaglenastat was dissolved in a solution of 25% (w/v) hydroxypropyl-β-cyclodextrin (HPBCD) in 10 mmol/L citrate, pH 2, prepared freshly before use. Etoposide (20 mg/mL; Accord) and cisplatin (1 mg/mL; Accord) were obtained from the pharmacy of Institut Gustave Roussy and handled according to the manufacturer’s instructions.

### Screening on inhibition of interaction between HIS-AIF_103-613_ and GST-CHCHD4

HIS-AIF_103-613_ and GST-CHCHD4 were produced in BL21+DE3(RIPL) bacteria during a 3h-induction with 0.5 mM IPTG at 37°C, as previously described ^11^. Assays were performed in white 384-well microplates (Greiner, #781075) with a total volume of 25 µL per well. Proteins, peptide N27 (positive control) ^11^ and beads were diluted in 0.1 % bovine serum albumin (BSA)/PBS, on ice.

A library containing in total 1.280 off-patent small molecules provided by Prestwick Chemicals was screened at 10 µM in 0.1% DMSO in monoplicate. First, 5 μL of 10 µM compound or 0.1 % DMSO / 3 µM N27 or 0.1 % DMSO / 0.1 % BSA PBS (negative control) was added. Secondly, 5 µL of His-AIF_103-613_ at 300 nM were incubated with compounds for 20 min before addition of 5 µL CHCHD4-Gst at 100 nM for 2 h. Thirdly, 10 µL of a mixture of 5 µg/mL donor and acceptors beads was added for 3 h at 23°C in the dark until reading on EnSpire® multimode plate reader (Perkin Elmer) (WO/2021/170963). For confirmation, best hits were validated in triplicate at concentrations of 10 µM and 30 µM.

### Computational study

The human AIF monomer three-dimensional structure (PDB ID: 1M6I) ^12^ cocrystallized with one flavin adenine dinucleotide (FAD) was used for docking calculations. Protein loop missing residues were added using SWISS-MODEL webserver ^13^. AIF structure was previously relaxed in explicit solvent (with 150 mM NaCl) by a molecular dynamics (MD) simulation in a cubic box with side 9.3 nm, using GROMACS software ^14^ with AMBER99SB-ILDN ^15^ and TIP3P ^16^ all-atom force fields. A 2-fs integration timestep, and LINCS constraints for h-bonds, were used. Short range cutoff for Coulomb and Van der Waals was set to 1.0 nm and particle mesh Ewald was used for long range electrostatics. The system was equilibrated with two short MD simulations (1 ns each), the first one in the NVT ensemble at 300 K using the V-rescale thermostat (with time constant 0.1 ps), the second one in the NPT ensemble at 1 bar using the Parinello-Raman barostat (with time constant 2.0 ps and compressibility 4.5 10-5 bar-1). Then, the system was submitted to a 500 ns production run. The 5 most populated clusters of the protein conformational ensemble were extracted from the MD trajectory and used as receptors in subsequent docking calculations. mitoxantrone was blindly docked 100 times on the whole surface of each of the AIF clusters (without specifying any preferred binding site), using AutoDock Vina ^17^. Since 10 binding modes are generated for each docking calculation, a total of 5,000 ligand binding modes were obtained for the AIF protein.

### Kaplan-Meier overall survival analysis by R2: Genomics Analysis and Visualization Platform

Datasets of patients with osteosarcoma (GSE42352), Ewing sarcoma (GSE17679), and rhabdomyosarcoma ^18^ were used to generate Kaplan-Meier curves for *AIFM1* gene or *CHCHD4* gene, comparing low expression to high expression groups, on the R2 database: Genomics Analysis and Visualization Platform (https://hgserver1.amc.nl/cgi-bin/r2/main.cgi). Expression cut-off was automatically calculated by R2’s algorithm, and Kaplan-Meier analysis was presented in graphs for each gene and each dataset.

### Cell lines and patient-derived xenografts (PDXs) cultures

Human pediatric osteosarcoma cell lines HOS parental (HOS), HOS-doxorubicin resistant (HOS-R/DOXO), and HOS-methotrexate resistant (HOS-R/MTX), MG63, U2OS, 143B, SAOS2, SAOS2-methotrexate resistant (SAOS2-R/MTX) and U2OS-Luc/mKate2 were routinely cultured in Dulbecco’s modified Eagle medium (DMEM), high glucose, GlutaMAX™ Supplement, pyruvate (Gibco, #31966021) supplemented with 10% FBS (Sigma-Aldrich, #F7524) and 1 % penicillin-streptomycin (Gibco, #15140122) ^19^. The PDXs GR-OS-10, GR-OS-18, GR-OS-12, GR-OS-15, GR-OS-9, GR-OS-11, GR-OS-20 were established from relapsed or refractory osteosarcoma tumor fragment (MAPPYACTS PDX project), secondary PDX cultures were performed by dissociating mechanically and enzymatically xenografted tumors and cultured in DMEM supplemented with 20% FBS ^20^. Cell lines and PDXs cells were incubated at 5 % CO2 and 37°C under mycoplasma-free conditions.

### Lactate Dehydrogenase (LDH) Assay

The half maximal inhibitory concentration (IC_50_) at 72h of all compounds for all cell lines was determined by Cytotoxicity Detection Kit ^PLUS^ (LDH), following the manufacturer’s protocol (Roche, #4744926001). This assay measures the conversion of a tetrazolium (INT) salt to a formazan product, which presents a reddish color. The intensity of color produced is proportional to the number of cells lysed. Absorbance was measured using a 96-well microplate reader at 490nm wavelength (TECAN Infinite M200). The IC_50_ was determined using [Inhibitor] vs. response --Variable slope (four parameters): Y=Bottom + (Top-Bottom)/(1+(IC_50_/X)^HillSlope) by GraphPad Prism 10.0 software (GraphPad Software Inc., California, USA).

### Clonogenic assay

Osteosarcoma cells were seeded at 500 cells/well in 6-well plates. After 24h, cells were exposed to mitoxantrone for different time durations (30 min, 1h, 2h, 4h, and 6h), then kept in the incubator for 10 days under standard conditions. On day 10 after the treatment, colonies were washed twice with PBS and incubated with Crystal Violet (Sigma-Aldrich, #HT90132) for 30 mins. Images were taken using Amersham™ ImageQuant™ 800 imaging system.

### siRNA transfection

To knock down AIF expression for evaluating the specificity of mitoxantrone, HOS cells were seeded at densities of 12.500, 18.750, 25.000, 31.250 cells/cm ^3^ and transfected with ON-TARGETplus Human AIFM1 siRNA (Dharmacon, #L-011912-00-0005) or ON-TARGETplus GAPDH Control Pool (Dharmacon, #D-001830-10-05) for 24h, 48h, 72h and 96h, using 0.2µL, 0.16µL, 0.2µL and 0.08µL of DharmaFECT 1 Transfection Reagent (Dharmacon, #T-2001-03), respectively, according to the manufacturer’s instructions. For subsequent experiments, a 48h time point was used. siRNA pool targeting AIF, comprising 4 selected siRNA duplexes each, has a modification pattern that addresses off-target effects caused by both strands (SMART pool).

### Co-immunoprecipitation

Eight million HOS cells were plated in 145 cm^2^ Petri dishes (Thermo Fisher, #168381) one day prior to treatment. Cells were treated with 7nM mitoxantrone during 48h or 0.0005 % DMSO (control). All cells were collected using a cell scraper, and lysates were prepared in 1 % NP-40 lysis buffer (50 mM Tris, pH 8.0, 150 mM NaCl, 1 mM MgCl_2_, 1 mM CaCl_2_, 10 % glycerol, phosphatase inhibitor (Roche, #4906845001), protease inhibitor cocktail (Roche, #05892970001)). Lysates were incubated on ice for 30 min and centrifuged at 4°C at maximum speed for 10 min. For immunoprecipitation, magnetic Dynabeads M-280 (Invitrogen, #11203D) were saturated with 1 % BSA for 1h to reduce nonspecific binding. Two micrograms of primary antibodies (anti-rabbit IgG control, Proteintech #30000-0-AP; anti-CHCHD4, 1:1000, Proteintech, #21090-1-AP) diluted in PBS with 0.5% Tween-20 (PBS-T) were added to the beads and incubated at room temperature with rotation for 20 min. Antibody-bead complexes were washed once with PBS-T, and protein samples were added, followed by overnight incubation at 4°C with rotation. Twenty-five micrograms of protein lysates were reserved as input. Antigen-antibody-bead complexes were washed twice with PBS-T, denatured, and reduced by boiling for 10 min in NuPAGE LDS sample buffer 4X (Thermo Scientific, #NP0008) containing dithiothreitol (DTT). Western blot analysis was performed as described, using antibodies against CHCHD4 (1:1000, Proteintech, #21090-1-AP), AIF (1:1000, Cell Signaling, #5318S), and goat anti-rabbit secondary antibodies (1:5000, Jackson ImmunoResearch, #111-035-144). Bands were visualized using Substrat HRP Immobilon Western (Millipore, #WBKLS0500) and imaged with an Amersham™ ImageQuant™ 800.

### Western blot

Immunoblotting was performed on cell lysates in nuclear and cytoplasmic extraction (NETN) buffer supplemented with phosphatase inhibitor (Roche, #4906845001) and protease inhibitor cocktail (Roche, #05892970001). Briefly, the cell lysates were separated on 10–15% Tris-glycine SDS–PAGE gel (Invitrogen, #NP0322) and transferred onto PVDF membrane. Membranes were blocked in 5% milk in TBS with 0.1% TWEEN-20 (TBST) for 1⍰hour at room temperature followed by overnight incubation with indicated primary antibodies (indicated in Table I) diluted in TBST containing 5% BSA. Membranes were washed and incubated with secondary antibody (Horseradish peroxidase (HRP)-labeled rabbit anti-mouse (1:5000, Jackson ImmunoResearch, #315-035-003) and goat anti-rabbit antibodies (1:5000, Jackson ImmunoResearch, #111-035-144) for 2 h at room temperature. Bands were visualized using the Substrat HRP Immobilon Western (Millipore, #WBKLS0500) and imaged with Amersham™ ImageQuant™ 800. Equal loading was verified by immunoblotting with Vinculin, GAPDH, then normalization of results was performed using free ImageJ software.

**Table I.** Primary antibodies used in Western blot. Primary antibodies were diluted in TBST, 5% BSA at 1:1000.

| Antibody Name | Reference |
| --- | --- |
| MICU1 | ATLAS Antibodies, #HPA037479 |
| COX17 | Atlas Antibodies, #HPA048158 |
| AIF | Cell Signaling, #5318S |
| CHCHD4 | Proteintech, #21090-1-AP |
| Total human OXPHOS cocktail | Abcam, #ab110411 |
| GLS1 | Proteintech, #12855-1-AP |
| Vinculin | Abcam, #ab219649 |
| VDAC | Cell Signaling, #4661 |
| phospho-Histone H2A.X (Ser139) | Active Motif, #AB_2793161 |

### Transmission electron microscopy

For ultrastructural studies, cells were fixed in 2 % glutaraldehyde in 0.1⍰M Sörensen phosphate buffer (pH 7.3) for 1h at 4°C, post-fixed with 2 % osmium tetroxide for 1h at room temperature and stained en bloc in 2 % uranyl acetate in 30 % methanol for 1h. Following dehydration through a graded ethanol series, cells were embedded in Epon™ 812. Polymerization was complete after 48⍰h at 60⍰°C. Ultrathin sections were stained with standard uranyl acetate and lead citrate and observed with FEI Tecnai 12 electron microscope. Digital images were taken with a SIS MegaviewIII CCD camera.

### ATP Assay

ATPlite 1 step Luminescence Assay system from PerkinElmer (#6016736) was used to evaluate the total ATP contents that reflect the catabolic/anabolic status of the cells by measuring total ATP concentration based on the reaction between ATP with added luciferase and D-Luciferine to produce Oxiluciferine and detectable light. Cells were cultured on white 96-well plates, provided with the kit (PerkinElmer) and total ATP content was measured 24h later with TECAN Infinite 200 Pro microplate reader.

### Bioenergetic profiling using Seahorse⍰XFe96 Assay

To determine the metabolic changes in cellular mitochondrial respiration, represented by Oxygen consumption rate (OCR), mitoxantrone-treated osteosarcoma cells were measured using the Seahorse XFe96 Analyzer (Agilent Technologies) in real-time in 96-well plates. For the untreated conditions, 2.000 cells/well were plated 72h before assay. In the treated conditions, 3000 cells/well, 4000 cells/well and 7000 cells/well were treated respectively during 24, 48, and 72h before analysis. Three different protocols were used to evaluate together or separately OCR and GlycoPER values by injecting different treatments, following manufacturer protocols; Seahorse XF Cell Mito Stress Test (Agilent Technologies, #103015-100); Seahorse XF Glycolytic Rate Test (Agilent Technologies, #103344-100); Seahorse XF Long Chain Fatty Acid Oxidation Stress Test Kit (Agilent Technologies, #103672-100)]. To count the cells and normalize the Seahorse analysis, cells were incubated with 2.5 μg/ml of Hoechst dye in 150 μL of PBS solution for 15 min at 37 °C and visualized and counted using Cytation 1 imaging system and Gen5 software (Agilent). Images were captured by DAPI overlay channels or phase-contrast. Raw data analysis and normalization were made using Seahorse analytics (Agilent).

### Mitochondrial membrane potential

HOS cells were seeded at 110.000 cells/3.5 cm ^2^ and allowed to adhere for 24h, before treatment of mitoxantrone at IC_50_ for 24h. Next, cells were trypsinized and incubated with 100 nM Tetramethylrhodamine methyl ester (TMRM) for 10 min at 37°C, in the dark. Data acquisition was performed using BD Accuri™ C6 Plus Flow Cytometer. At least 10.000 events were acquired for each sample, and all data were analyzed using FlowJo™ Software (BD Life Sciences).

### Ca^2+^ measurement

Osteosarcoma cells were seeded onto 35 × 15 mm Petri dishes and incubated overnight. Following treatment with the IC_50_ of mitoxantrone for 24h or under basal conditions, cells were washed twice with modified Krebs-Ringer buffer (KRB) containing 135 mM NaCl, 5 mM KCl (Sigma, #P9541-500G), 1 mM MgCl_2_·6H_2_O (Euromedex, #2189-C), 0.4 mM KH_2_PO_4_ (Euromedex, #2018), 1 mM MgSO_4_·7H_2_O (Euromedex, #P027-A), and 20 mM HEPES (Gibco, #15630-056). Cells were then incubated at 37 °C for 30 min in supplemented (5.5 mM glucose and 100 µM EGTA) KRB containing 4 µM Rhod-2 AM (Invitrogen, #R1245MP) and 25 µg/mL Hoechst 33342 (Sigma, #14533-100MG). After staining, cells were washed twice with KRB and maintained in 2 mL of supplemented KRB (with glucose and EGTA) before measurement. Basal fluorescence intensity of Rhod-2 was recorded for 5 min at 2 s intervals using a Cytation 1 imaging reader (Agilent) with 488- and 561-nm excitation filters and a 4× objective. Regions of interest were selected randomly. After baseline recording, 250 µL of KRB solution containing 100 µM histamine and 200 mM CaCl_2_ was added, and fluorescence changes were recorded for an additional 30 min (interval = 2 s). Images were analyzed using Gen5 software (Agilent). Mean fluorescence intensity was calculated at each time point by dividing the total Rhod-2 signal by the number of nuclei stained with Hoechst 33342 in the corresponding region of interest.

### Metabolomics

#### a, Sample preparation

HOS (300.000 cells/well) were cultured in 6 well-plates and treated with IC_50_ of mitoxantrone during 72h. Cells were gently rinsed with cold PBS and lysed by adding 500 µL of cold methanol/water (9/1, v/v,-20°C) with internal standards (ISTD). After a quick scrap, supernatants were pooled, with two wells per condition in a single microtube. Supernatants in microtubes were vortexed at 800 g for 5 min and centrifuged at 15000⍰g for 10⍰min at 4⍰°C. Supernatants were split in two parts: 150 µL were used for GC-MS experiment in injection vial and 300 µL were used for UHPLC-MS experimentation. LC-MS aliquots were evaporated and dried extracts were resuspended with 150 µL of MilliQ water. Samples were kept at-80°C until injection or transferred in vials for direct analysis by UHPLC/MS. After evaporation, dry GC-MS aliquot was spiked with 50 µL of methoxyamine (20 mg/mL in pyridine) and stored at room temperature in the dark overnight. Then, 80 µL of N-Methyl-N-trimethylsilylfluoroacetamide (MSTFA) was added and final derivatization occurred at 40°C for 30 min. Samples were directly injected into GC-MS.

#### b, Targeted analysis of nucleotides and cofactors by ion pairing ultra-high performance liquid chromatography (UHPLC) coupled to a Triple Quadrupole (QQQ) mass spectrometer

Targeted analysis was performed on a RRLC 1290 system (Agilent Technologies) coupled to a Triple Quadrupole 6470 (Agilent Technologies) equipped with an electrospray source. Ten μL of sample were injected on a Column Zorbax Eclipse XDB-C18 (100 mm × 2.1 mm particle size 1.8 µm) from Agilent technologies. Gradient mobile phase consisted of water with 2mM of dibutylamine acetate concentrate (DBAA) (phase A) and acetonitrile (phase B). Flow rate was set to 0.4 mL/min, and gradient was performed as follows: initial condition was 90 % phase A and 20 % phase B, maintained during 3 min. Molecules were then eluted using a gradient from 10% to 95 % phase B over 1 min. The column was washed using 95 % mobile phase B for 2 min and equilibrated using 10 % mobile phase B for 1 min. Scan mode used was the MRM for biological samples. Peak detection and integration of the analytes were performed using the Agilent Mass Hunter quantitative software (B.10.1).

#### c, Widely-targeted analysis of intracellular metabolites gas chromatography (GC) coupled to a triple quadrupole (QQQ) mass spectrometer

GC-MS/MS method was performed on a coupling gas chromatography / triple quadrupole 7890B / 7000C (Agilent Technologies, Waldbronn, Germany). The scan mode used was the MRM for biological samples. Peak detection and integration of analytes were performed using the Agilent Mass Hunter quantitative software (B.07.01), exported as tables and processed with R software (version 4.0.3) and the GRMeta package (Github/kroemerlab).

#### d, Data analysis

Data analysis was conducted using the Statistical Analysis module and Enrichment Analysis model by MetaboAnalyst 6.0 platform. Log transformation (base 10) was used to transform and normalize raw data.

#### Measurement of intracellular glutamate/glutamine ratio

HOS and U2OS were seeded in 96-well plates at 2.000 cells/well and treated with mitoxantrone at IC_25_ for 72h. Semi-quantitative analysis of intracellular glutamate and glutamine levels was performed using the Glutamine/glutamate-Glo™ Assay (Promega, #J8021) following the manufacturer’s protocol. Briefly, glutaminase solution or enzyme-free buffer was added to cell lysates for the detection of total glutamine and glutamate or glutamate-only levels, respectively. All samples were mixed with glutamate detection reagent containing luciferin and the luminescence was recorded after 60 min of incubation. All values were normalized to cell numbers in a replica plate using Hoechst staining in PBS (1:4000) and automated counting in BioTek Cytation 1 (Agilent). To obtain the glutamine signal, the glutamate-only signal was subtracted from the total glutamine and glutamate signal. The glutamate/glutamine ratio was calculated and compared between untreated and mitoxantrone-treated cells.

#### SynergyFinder

Evaluation of drug interactions between compounds was conducted on U2OS, 143B, SAOS2, HOS models. HOS, U2OS, 143B (2000 cells/well), and SAOS2 (5000 cells/well) were seeded in 96-well plates and treated with a combination of mitoxantrone and telaglenastat, in the form of a matrix, for 72h. Cell viability was measured using LDH assay, and data was uploaded on the SynergyFinder platform (https://synergyfinder.fimm.fi/). Using the highest single agent (HSA) model, an overall synergy score was calculated for each matrix, with values below-10 indicating antagonism, in between-10 and 10 as additivity, and above 10 as synergism. 2D and 3D interaction maps were also provided, in which the colors green, white, and red demonstrate antagonism, additivity, and synergism, respectively. Experiments were repeated at least 3 times.

#### Nucleoside rescue assay

HOS and U2OS (2.000 cells/well) were seeded and treated with 1x (30 μM each of cytidine, guanosine, uridine, adenosine and 8 μM thymidine) or 2x (60 μM each of cytidine, guanosine, uridine, adenosine and 16 μM thymidine) dilution of EmbryoMax® Nucleosides 100x (Sigma-Aldrich, Cat#ES-008-D) in a medium containing mitoxantrone and CB-839 at IC_25_ doses for 72h. Cell viability was evaluated using the LDH assay.

#### Animal studies

All *in vivo* experiments were conducted following the approval of the CEEA26 Ethic Committee (approval number: APAFIS #37810-2022062308201583) agreed by the French Ministry of Research regulations (articles R.214-87 à R.214-126). Animals were obtained and kept following standard animal regulations, health and care, and ethical controls in Gustave Roussy animal facility (Villejuif, France). A total of 8 × 10^6^ U2OS-Luc/mKate2-cells were injected subcutaneously into thirty six female immunocompromised NOD.Cg-Prkdc^scid^ IL2rg^tm1Wjl^/SzJ (NSG) mice. Local anesthesia using lidocaine was used for subcutaneous injection. Treatment started at advanced tumor stage, when tumors reached 60–100 mm^3^ about four weeks after engraftment. Mice bearing established tumors were blindly randomized into four groups: control group (PBS + HPBCD) or mitoxantrone monotherapy group (mitoxantrone + HPBCD) or telaglenastat monotherapy group (PBS + telaglenastat) or combination group (mitoxantrone + telaglenastat). Treatments were administered as follows: telaglenastat or HPBCD was given orally five days/week for three weeks, and mitoxantrone or PBS was administered intraperitoneally 2 doses on the first and fourth day. Clinical observations such as skin changes, behavior, posture, responses to handling, and abnormal movements were registered. Body weight and tumor volume were measured twice weekly. Tumor volume was calculated using the equation: V (mm3) =width2 (mm2) ×length (mm)/2. The experiments lasted until tumors reached specific endpoints (maximum 1500 mm^3^), as detailed in the ethical projects.

### Generation of *MICU1* knocked-out HOS clones

#### a, gRNA design, cloning and testing

All guide RNA (gRNA) binding sites were generated from CRISPOR gRNA design online tool (http://crispor.tefor.net/) to target intron 2 and intron 16 of *MICU1* gene and were inserted into the phU6 gRNA plasmids (Addgene #53188). gRNA sequences are 5’-CACCGCTCTACCAAGTCAGACTAG-3’ to target intron 2 and 5’CACCACAGGGGCCCAGCCTCATAT-3’ to target intron 16. The dephosphorylated digested plasmid backbones were ligated with the phosphorylated annealed gRNA to generate plasmid containing gRNA. Plasmid containing gRNA and Cas9 plasmids (Addgene #57818) were transformed into *E. coli* competent cells via heat shock and were then clonally expanded for plasmid extraction (Machery-Nagel NucleoBond Xtra Midi or Maxi kit), as previously described ^21^. gRNA efficiency was tested following the procedures provided in ENIT protocols, as previously described ^22^.

#### b, Plasmid transfection

Briefly, 150.000 cells were seeded into 6-well plates and transfected the next day (70-80 % confluence) with 2 μg total plasmid, 0.5 μg of each plasmid containing gRNA and 1μg p-CMV-Cas9-GFP. After 48h post-transfection, GFP-positive cells were sorted by Accuri™ C6 Flow Cytometer (BD Biosciences) to separate one cell per well in 96-wells plates. Clone validation was ensured with different methods: PCR, DNA sequencing and Western Blot of protein expression

### Generation of MICU1 overexpressed HOS clones

#### a, Plasmid and transfection

HOS cells were seeded at 1.5 × 10^5^ cells/well in 6-well plates 24h before transfection. The pLYS1-MICU1-Flag plasmid (Addgene #50058) was transfected using TurboFect (Thermo Scientific, #15325016) following the manufacturer’s instructions.

#### b, Antibiotic selection and single-cell cloning

24h after transfection, cells were placed under puromycin (0.2 µg/mL) selection for 5 days to eliminate non-transfected and transiently transfected cells. Surviving cells were then individually sorted on each well of 96-well plate using a BD FACSAria™ Fusion UV (BD Biosciences).

#### c, Clone validation

After expanding, PCR was performed to confirm MICU1-Flag integration using a forward primer within the MICU1 gene body (5⍰-CTTGTCATCGTCATCCTTGTAATC-3⍰) and a reverse primer targeting the FLAG-tag sequence (5⍰-GCCATGCAGAGGCAGCTCAAGAAG-3⍰). Amplicons were analyzed by agarose gel electrophoresis under standard conditions. MICU1 overexpression was verified by Western blotting using antibodies against MICU1 (ATLAS Antibodies, HPA037479).

#### Statistics

Statistical analysis was performed using GraphPad Prism 10.0. One-way ANOVA test was used for statistical analysis of biological experiments: p<0.05 (*), p<0.01 (**), p<0.001 (***), p<0.0001 (****). Two-way ANOVA (mixed model) was used for tumor volume curves: p > 0.1234 (ns - not significant); p < 0.0332 (*);p < 0.0021 (**); p < 0.0002 (***);p < 0.0001 (****).

## Results

### High-throughput screening identifies mitoxantrone as an AIF/CHCHD4 interaction inhibitor

Given the essential role of the AIF/CHCHD4 complex in mETC biogenesis and mitochondrial function ^11^, we developed a robotized high-throughput screening (HTS) assay using the Alphascreen technology to discover compounds capable of disrupting this interaction in an anti-cancer therapeutic perspective. For this assay, recombinant HIS-tagged AIF_103-613_ and GST-tagged CHCHD4 proteins were purified and coupled to the corresponding donor and receptor alpha beads, as previously described ^8^. After the optimization of protein and beads concentrations and incubation times, the interaction between the two proteins was quantified by light emission. A synthetic peptide, named N27, corresponding to the N-terminal 27 amino acids of CHCHD4, served as positive control to fully disrupt the AIF/CHCHD4 interaction. After validating assay robustness (Z’ factor 0.5–1), we screened the Prestwick library of 1.280 FDA-approved molecules. Compounds (tested at 10 µM and 30 µM) were evaluated for their ability to reduce the alpha signal, relative to the positive control N27. From this screen, we identified 148 out of 1280 candidate hits that disrupted the AIF/CHCHD4 complex, including several clinically repositionable molecules (Fig. 1a). Eleven compounds were prioritized for further validation according to two criteria: (i) the signal at 10 μM was lower than the threshold value [TH = mean (negative control signal) – 5 * SD (negative control signal)], and (ii) the signal at 30 μM was lower than that at 10 μM (Fig Supp. 1a). Compounds were considered as AIF/CHCHD4 inhibitors if they achieved at least 50% inhibition of the interaction, compared to the N27 positive control.

**Figure 1.**
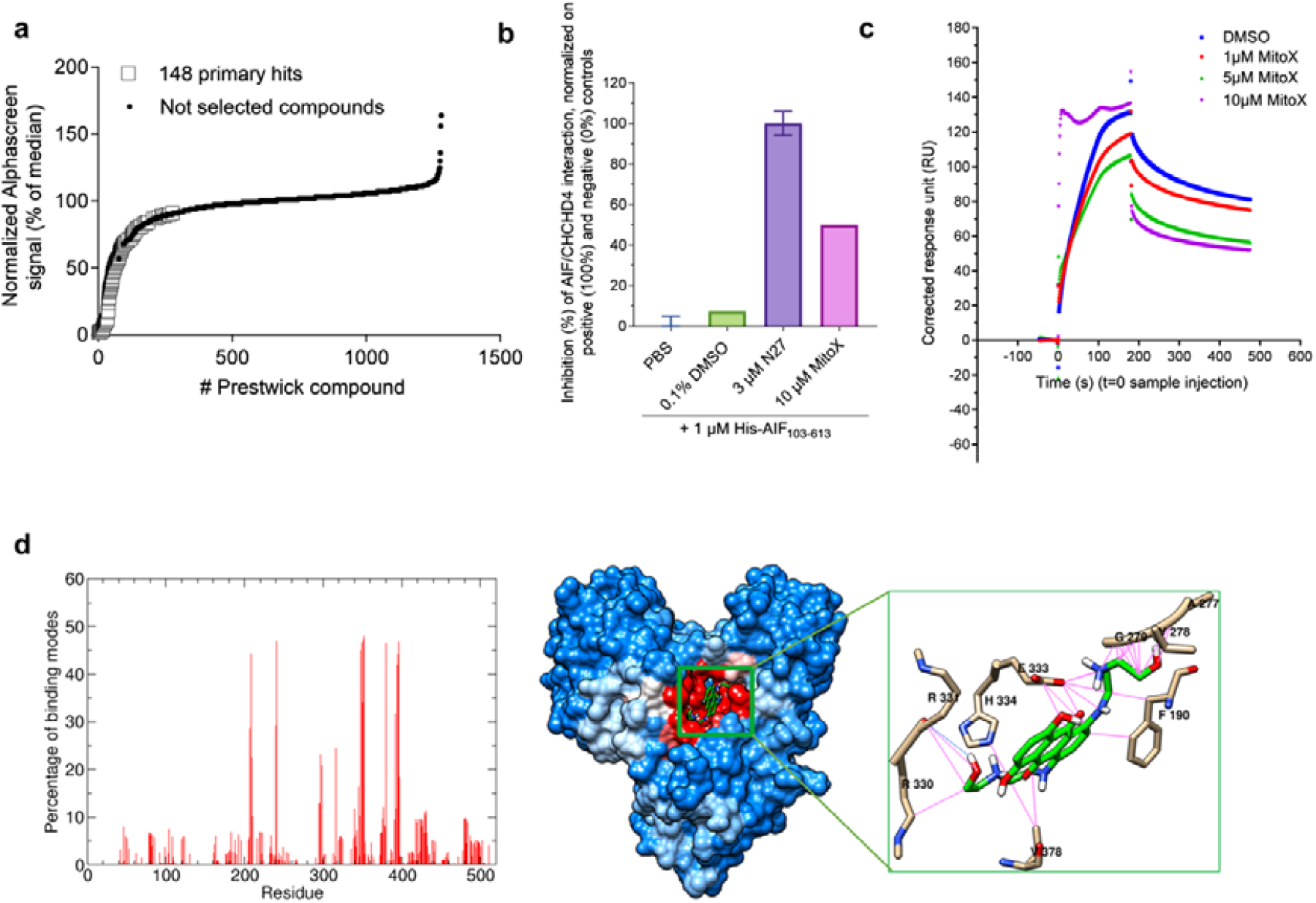
High-throughput screening identification of mitoxantrone as disruptor of the AIF/CHCHD4 complex. a, Inhibition of HIS-AIF_103-613_ / GST-CHCHD4 interaction. Primary screening of 1280 Prestwick compounds at 10 µM by Alphascreen, ordered by percentage of median. Primary hits (148) are represented by write squares; black dots are not selected compounds. For each of the four 384-well-plates, the threshold (TH) was set at: TH = mean *(negative control)* – 5 * SD *(negative control)*. Every raw signal < TH was selected as a primary hit. The screening was processed once in monoplicate. b, Surface plasmon resonance (SPR) counterscreen of primary hits from Alpha screening. 10 µM mitoxantrone were incubated during 20 min with 1 µM AIF in pH 7.4 DPBS at RT. Then the mixture was injected during 3 min, 30 µL per min, on immobilized GST-CHCHD4. 0.1 % DMSO / 1 µM AIF was a negative control. 3 µM N27 peptide was a positive control. c, SPR sensorgrams for HIS-AIF_103-613_ binding to immobilized GST-CHCHD4 in presence of mitoxantrone. mitoxantrone was tested at increasing concentrations from 0.5 µM to 10 µM. d, Left panel, Contact frequency of AIF residues with the 5,000 mitoxantrone binding modes generated by docking calculations; right panel, localization of AIF residues most frequently in contact with mitoxantrone (blue, white, and red colors indicate residues that are contacted less than 10 %, between 10 % and 20 %, and more than 20 % out of all the ligand binding modes, respectively), and focus on AIF residues that interact with mitoxantrone in the best docking binding mode complex.

Among the prioritized eleven hits, mitoxantrone was selected for surface plasmon resonance (SPR) experiments based on its known therapeutic activity in cancer for intercalating into DNA and inducing DNA strand breaks (Fig Supp. 1b). Subsequently, SPR experiment confirmed that mitoxantrone inhibited the AIF/CHCHD4 interaction by 50 % at 10 µM and decreased resonance was observed while increasing mitoxantrone concentration from 1 to 10 µM, in line with AlphaScreen assay above (Fig. 1b, c). Then, mitoxantrone was blindly docked on AIF monomer *in silico* using the protocol described in the Material and methods section. Its preferred binding site on AIF was examined by calculating the frequency of contact between AIF residues and the 5,000 binding modes generated by docking (Fig. 1d). These data clearly demonstrate that mitoxantrone binds preferentially to the NADH pocket. In the best binding mode of mitoxantrone to AIF, the complex is stabilised by hydrophobic interactions between the anthraquinone moiety and the protein residues Phe190 and Val378. It is also stabilised by electrostatic interactions of the two hydrophilic side chains of mitoxantrone with protein V278, G279, and E333 residues on the one hand, as well as with R330, R331, and H334 residues on the the hand (Fig. 1d). This strongly suggests that mitoxantrone competitively inhibited NAD binding, which in turn would prevent AIF dimerization.

### Cytotoxic effects of mitoxantrone in osteosarcoma models

To select the preclinical models to perform further functional experiments of disrupting the AIF/CHCHD4 complex, we first explored the *AIFM1* and *CHCHD4* mRNA expression in three pediatric sarcomas: osteosarcoma, Ewing’s sarcoma and rhabdomyosarcoma, using publicly available data on the R2 platform. Expression cut-off was computed automatically by the algorithm of the R2 platform to divide patients into 2 groups of low/high expression of *AIFM1* and *CHCHD4* mRNA expression for Kaplan-Meier analysis with a log-rank statistical test. Patients with high *CHCHD4* mRNA expression showed a lower overall survival rate than patients with low *CHCHD4* mRNA expression for all three sarcoma types (osteosarcoma: p = 0.024; Ewing’s sarcoma: p = 0.017, rhabdomyosarcoma: p = 0.107) (Fig. 2a). For *AIFM1* mRNA, high expression was associated with lower overall survival in patients with osteosarcoma (p = 3.84e-04) and Ewing sarcoma (p = 4.12e-05). Rhabdomyosarcoma cases with highly expressed *AIFM1* mRNA had a better overall survival compared to low-expressed *AIFM1* (p = 0.119) (Fig. 2b). In addition, our previous analysis of the RNA-seq transcriptomic results of patient cohorts from MAPPYACTS and OS2006 showed significant levels of *AIFM1* and *CHCHD4*, for both newly diagnosed and relapsed osteosarcoma ^23 24 8^. Considering these results, osteosarcoma was chosen for our studies in preclinical models.

**Figure 2.**
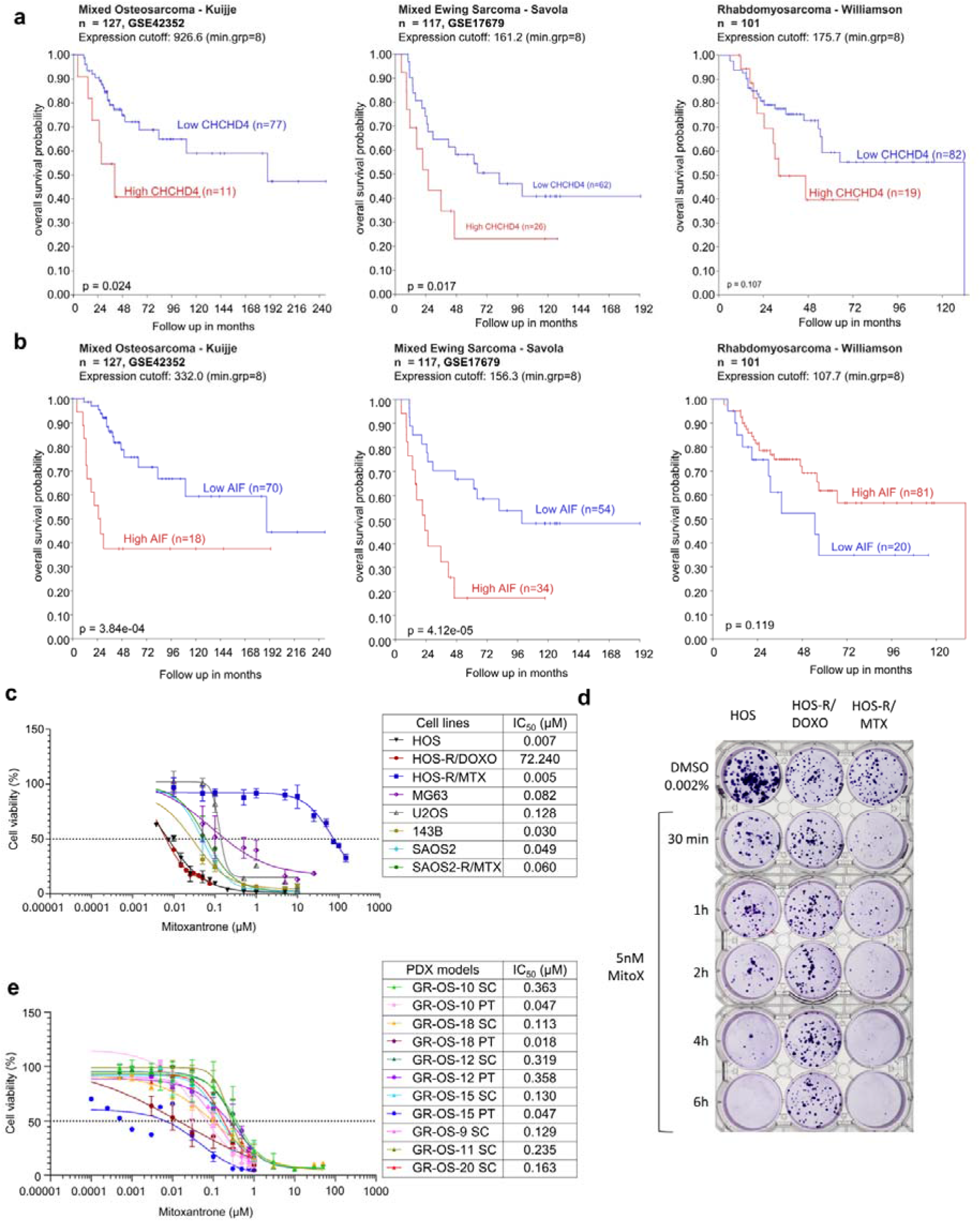
Mitoxantrone efficiently exhibits cytotoxicity on different osteosarcoma models. **a, b**, Kaplan-Meier overall survival analysis for patients with osteosarcoma, Ewing sarcoma and in the GSE42352, GSE17679 dataset, respectively. Rhabdomyosarcoma data set was collected from Missiaglia et al., 2012 ^18^. Kaplan-Meier curves for a, *AIFM1* gene and b, CHCHD4 gene of low-expression versus high-expression groups based on median AIFM1 and CHCHD4 expression, respectively. Kaplan-Meier analysis and automatically computed expression cut-off were generated by R2: Genomics Analysis and Visualization Platform. **c**, Cell viability of 8 pediatric osteosarcoma cell lines treated with mitoxantrone. Dose-response curves represent cell viability measured using a LDH assay at increasing concentrations of mitoxantrone after 72h of treatment. Individual experiments are shown (n > 3). Data were shown as mean ± SD. IC_50_ inhibitory concentration yielding 50% cell viability. 0.0005% DMSO was used as the solvent of mitoxantrone and as control for calculation of percentage of viability. **d**, Representative image of a 6-well plate showing fixed colonies subjected to crystal violet staining of HOS, HOS-R/DOXO, and HOS-R/MTX cells after treatment with either 0.0005 % DMSO as negative control or mitoxantrone at 5nM (IC_50_ of HOS-R/MTX) during indicated time points. **e**, Cell viability of 11 secondary cultured *in vitro* PDX samples derived from refractory or relapsed tumors treated with different concentrations of mitoxantrone, was measured using an LDH assay. Dose-response curves represent cell viability at increasing concentrations of mitoxantrone after 72h of treatment and the IC_50_ values were determined using GraphPad Prism software. Individual replicates are shown (n > 3). Data were shown as mean ± SD.

To assess the cytotoxic activity of mitoxantrone, a panel of eight human osteosarcoma cell lines, including different parental and drug-resistant lines (HOS, HOS-R/DOXO, HOS-R/MTX), MG63, U2OS, 143B, SAOS2 and SAOS-R/MTX) were exposed to different concentration of mitoxantrone for 72h before measurement of cell viability using LDH assay. The IC_50_ values for the osteosarcoma cell lines ranged from 0.005 μM to 0.128 μM, except for HOS-R/DOXO at 72.250 μM and thus considered as resistant (Fig. 2c). Given that mitoxantrone is a doxorubicin structural analog, one possible explanation is that mitoxantrone might share the same resistance mechanism, mediated by increased P-glycoprotein efflux activity ^25 26^. In a clonogenic assay, parental and resistant HOS cells were exposed to mitoxantrone during indicated time points. and subsequently cultured in drug-free medium for the complete assay duration of 10 days. Parental HOS and HOS-R/MTX cells were highly sensitive to 5nM mitoxantrone (IC_50_ of HOS-R/MTX, the lowest IC_50_ among three cell lines). The inhibitory effect of mitoxantrone on HOS and HOS-R/MTX increased progressively with treatment duration, with an immediate reduction in colony-forming ability after 30 min of exposure, and no surviving clones detectable after only 6h of mitoxantrone treatment. In contrast, HOS-R/DOXO cells displayed strong resistance to mitoxantrone, with cell density remaining unchanged following exposure to 5 nM mitoxantrone for up to 6h (Fig. 2d) (Fig. 2d).

Next, eleven secondary PDX cultures were tested, which were obtained from paratibial and subcutaneous PDX models originated from seven patients with refractory and relapsed osteosarcoma within the MAPPYACTS trial ^20^. These models reflect the heterogeneity and complexity of osteosarcoma while maintaining its molecular features, genomic landscape, and patterns of sensitivity or resistance to chemotherapeutic agents observed in patient tumors ^20^. Secondary PDX cultures derived from subcutaneous and paratibial PDX were dissociated by mechanistical and enzymatic methods, cultured *in vitro*, and treated with mitoxantrone. The IC_50_ values of mitoxantrone for all PDX models ranged from 0.018 μM to 0.363 μM, indicating a comparable sub-micromolar efficacy to that observed in the cell line model panel (Fig. 2e). In addition, to compare the efficacy of mitoxantrone to standard chemotherapeutic agents, all PDXs and three osteosarcoma cell lines (HOS, HOS-R/DOXO and HOS-R/MTX) were treated with etoposide or cisplatin and showed sensitivity to these compounds at low to medium micromolar concentrations (IC_50_ = 0.261 - 9.854 µM and 2.192 - 3.478 µM for cell lines, IC_50_ = 6.84 - 120 µM and 5.33 - 34.91 µM for PDXs, respectively) (Fig Supp. 1b, c). Altogether, our data suggests that mitoxantrone preserves high efficacy in the parental cell line and its methotrexate or doxorubicin resistant cells, as well as in PDX culture models derived from refractory or relapsed osteosarcoma, within a concentration range of conventional chemotherapies used in primary osteosarcoma treatment protocols.

### Mitoxantrone disrupts the AIF/CHCHD4 complex and induces alterations in mitochondrial functions

Co-immunoprecipitation analysis demonstrated the disruption of the AIF/CHCHD4 complex in parental HOS cells treated by mitoxantrone after 48h (Fig. 3a). We next knocked down AIF by transfecting HOS cells with a siRNA pool against *AIFM1* and subjected them to mitoxantrone treatment. Importantly, mitoxantrone was four times less effective in *AIFM1*-knocked-down cells (IC_50_ = 0.363 μM), which indicates the relevance of AIF in mitoxantrone’s cytotoxicity in this cell line (Fig. 3b). As previously reported, the AIF/CHCHD4 complex constitutes the central disulfide relay system on the mitochondrial IMS, ensuring the oxidative folding and stability of cysteine-rich substrates such as MItochondrial Calcium (Ca^2+^) Uptake 1 (MICU1) (Ca^2+^ homeostasis) and Cytochrome C Oxidase Copper Chaperone COX17 (COX17, CIV) (mETC assembly), which are essential for mitochondrial function ^11^. Therefore, we next investigated the consequences of disrupting the AIF/CHCHD4 import machinery on the protein expression levels of these substrates in HOS cells. The results revealed a time-dependent depletion of AIF, CHCHD4, and their downstream substrates MICU1 and COX17 within 24 - 72h of treatment with the IC_50_ dose of mitoxantrone. Notably, the expression level of voltage-dependent anion channel (VDAC), a mitochondrial outer membrane protein independent of the AIF/CHCHD4 axis, remained unchanged, suggesting there was no alteration in terms of mitochondrial mass (Fig. 3c). Moreover, as the AIF/CHCHD4 disulfide-relay pathway also contributes to mETC biogenesis, particularly in Complex I (CI) and Complex IV (CIV) assembly ^9^, we decided to examine the protein expression of five mETC complexes. After 48h treatment with mitoxantrone, both cell lines demonstrated a significant downregulation in CI and CIV compared to the control (Fig. 3d). Together, these data indicate that mitoxantrone selectively disrupts the AIF/CHCHD4 import machinery without reducing mitochondrial content, thereby impairing the protein expression of their different substrates including MICU1, COX17, as well as mETC complexes CI and CIV, which is consistent with the prior description of AIF/CHCHD4-dependent respiratory chain assembly role ^11^.

**Figure 3.**
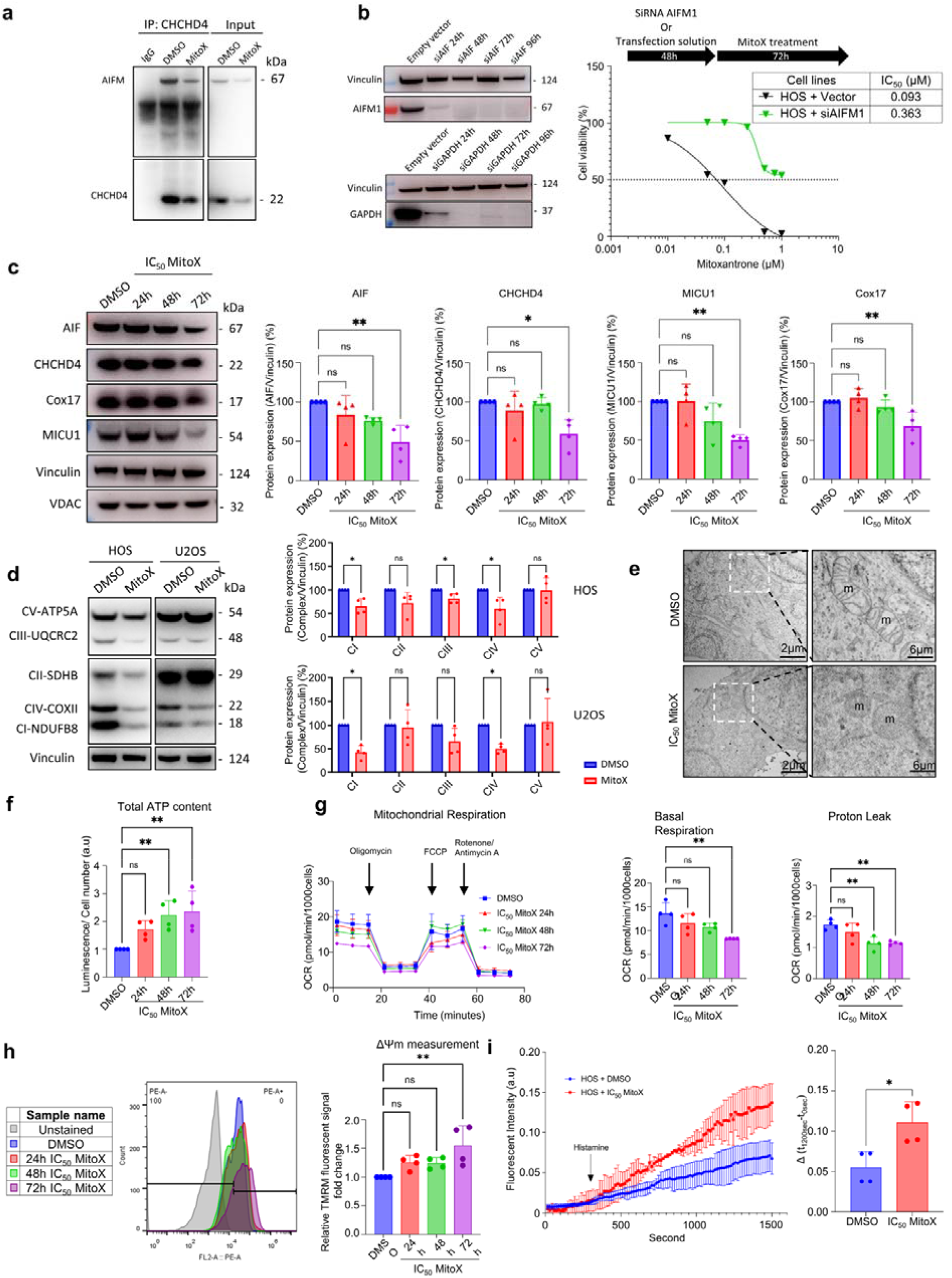
Mitoxantrone interrupts AIF/CHCHD4 interaction and perturbs energetic metabolism, mitochondria in osteosarcoma cell lines. **a**, AIF/CHCHD4 co-immunoprecipitation using CHCHD4 antibody to immunoprecipitate whole protein fraction from HOS cells treated with DMSO or with mitoxantrone during 48h. Immunoprecipitation was followed by Western blotting with AIF and CHCHD4 immunostaining. IgG immunoprecipitated sample was used as negative control. **b**, Representative Western blot analysis of AIF-siRNA (siAIF) treatment over time (left panel), with siGAPDH as positive control. Based on this, AIF was silenced 48h before exposure to increasing concentrations of mitoxantrone. Vinculin was used as loading control. Cell viability is measured using an LDH assay. Dose-response curves show cell viability at increasing mitoxantrone concentrations after 72h of treatment (n = 3). Data were shown as mean ± standard deviation (SD). **c**, Western blot analysis showing the expression levels of AIF, CHCHD4 and their substrates Cox17 and MICU1 in response to either 0.0005% DMSO as a negative control or IC_50_ of mitoxantrone treatment over time (6, 24, and 48h). Vinculin and VDAC were used as loading controls. Data are presented as the mean ± SD from four independent experiments (n = 4). Statistical significance was assessed using ANOVA with Sidak’s correction for multiple comparisons. Significance levels are indicated as follows: not significant (ns), p < 0.0332 (*), p < 0.0021 (**), p < 0.0002 (***), and p < 0.0001 (****). **d**, Western blot analysis of mitochondrial electron transport chain complexes in osteosarcoma cell lines HOS and U2OS treated with either 0.0005% DMSO (control) or IC_50_ mitoxantrone during 48h. Vinculin was used as loading control. **e**, Representative transmission electron microscopy images of HOS cells treated with either 0.0005% DMSO as negative control or with IC_50_ of mitoxantrone for 48h, showing that the compound induced cristolysis and changes in mitochondrial ultrastructure. m: mitochondrion **f**, ATP levels were semi-quantified over time in HOS cells pre-treated with either 0.0005% DMSO as a negative control or with IC_50_ mitoxantrone for 6, 24, and 48h, using the ATPlite assay. Data are presented as the mean ± SD from four independent experiments (n = 4). Statistical significance was assessed using ANOVA with Sidak’s correction for multiple comparisons. Significance levels are indicated as follows: not significant (ns), p < 0.0332 (*), p < 0.0021 (**), p < 0.0002 (***), and p < 0.0001 (****). **g**, Seahorse XFe96 Mito Stress Test graph displaying the oxygen consumption rate (OCR; pmol/min/1.000 cells) over time in HOS cells pre-treated with either 0.0005% DMSO as a negative control or with IC_50_ mitoxantrone for 24, 48, and 72h (left). The test was conducted using the following compounds: oligomycin (2.5 µM), Carbonyl cyanide-p-trifluoromethoxyphenylhydrazone (FCCP) (1 µM), and rotenone/antimycin A (0.5 µM). Basal respiration and proton leak values were extrapolated from the kinetic graph. OCR measurements were normalized to cell counts determined by nuclei DAPI staining. Data represent the mean ± SD of three independent experiments, each with at least six technical replicates. Statistical analysis was performed using ANOVA with Sidak’s correction for multiple comparisons, with significance levels indicated as follows: ns, not significant; p < 0.0332 (*); p < 0.0021 (**); p < 0.0002 (***);p < 0.0001 (****). **h**, Mitochondrial membrane potential assessed by flow cytometry using 100 nM TMRM fluorescent probe labelling for 20 min. HOS cells were analyzed at different time points following mitoxantrone treatment. Oligomycin and FCCP were used as positive controls for membrane potential modulation, and unstained cells served as negative controls for gating. Four independent experiments were performed (n = 4). Data represent the mean±SD. Statistical significance was assessed using ANOVA with Sidak’s multiple comparison test correction, with significance levels indicated as follows: ns, not significant; p < 0.0332 (*); p < 0.0021 (**); p < 0.0002 (***);p < 0.0001 (****). **i**, Histamine-induced mitochondrial Ca^2+^ uptake in osteosarcoma cells measured using the fluorescent probe Rhod-2 AM. Cells were treated with 0.0005% DMSO (control) or IC_50_ mitoxantrone for 24h, loaded with Rhod-2 AM (4 µM, 30 min) and Hoechst 33342 for nuclear counterstaining, and imaged using a Cytation 1 reader (Agilent). Baseline fluorescence was recorded for 5 min (interval = 2 s) before stimulation with histamine (100 µM) in the presence of extracellular CaCl_2_ (200 mM), and changes in Rhod-2 fluorescence were monitored for an additional 30 min. Mean fluorescence intensity was normalized to Hoechst-positive nuclei. Data are presented as the mean ± SD from four independent experiments (n = 4). Statistical significance was assessed using an unpaired two-tailed Student’s t-test. Significance levels are indicated as follows: not significant (ns), p < 0.0332 (*), p < 0.0021 (**), p < 0.0002 (***), and p < 0.0001 (****).

As the AIF/CHCHD4 complex orchestrates the correct import and folding of proteins contributing to various mitochondrial activities, we next assessed the impact of mitoxantrone treatment on mitochondrial morphology and functions. Transmission electron microscopy revealed substantial morphological alterations in the mitochondrial ultrastructure in mitoxantrone-treated HOS cells, as compared to controls. While mitochondria typically appeared elongated, with well-organized cristae and a clear matrix consistent with normal respiratory function, mitoxantrone-treated HOS cells displayed swollen and rounded mitochondria, characterized by a dense matrix and by cristae loss, indicative of potential cristolysis (Fig. 3e). We subsequently studied mitoxantrone-induced changes in mitochondrial bioenergetics. By measuring the total ATP content in HOS cells, we observed a time-dependent increase in total ATP levels upon treatment with mitoxantrone, compared to control cells (Fig. 3f). The Agilent Seahorse XF Mito Stress Test, which allows for real-time analysis of mitochondrial energetic functions, revealed that mitoxantrone significantly reduced oxygen consumption rate (OCR) in HOS cells at 72h post-treatment, accompanied with lower proton leak (Fig. 3g), indicating a possible energetic reprogramming after mitoxantrone treatment. Tetramethylrhodamine, methyl ester (TMRM) measurements of mitochondrial membrane potential (ΔΨm) showed progressive mitochondrial hyperpolarization after mitoxantrone exposure, suggesting an energetic reprogramming towards a high-potential, low-respiration state (Fig. 3h).

Furthermore, since the protein expression level of the AIF/CHCHD4 complex substrate MICU1 decreased upon mitoxantrone treatment and Ca^2+^ is a versatile signalling messenger that regulates vital as well as lethal cellular functions, we evaluated the effects of mitoxantrone on mitochondrial Ca^2+^ signaling. We developed a mitochondrial Ca^2+^ assay using Rhod-2 fluorescence dye, with histamine addition to induce endoplasmic reticulum Ca^2+^ release and subsequent mitochondrial Ca^2+^ uptake. Histamine stimulation resulted in approximately a two-fold higher increase in mitochondrial Ca^2+^ accumulation in mitoxantrone-treated HOS cells as compared to controls, suggesting that mitoxantrone enhanced mitochondrial Ca^2+^ uptake capacity (Fig. 3i).

To investigate if MICU1 misregulation was a downstream consequence of mitoxantrone-induced AIF/CHCHD4 binding interruption, we generated human *MICU1* knockout (HM1KO) and MICU1-overexpressing (HM1OE) HOS cell lines using CRISPR-Cas9 and stable plasmid transfection, respectively, and assessed their Ca^2+^ uptake capacity and other mitochondrial functions (Fig Sup. 2a). Mimicking mitoxantrone-treated cells, HM1KO cells exhibited a significant increase in mitochondrial Ca^2+^ uptake capacity upon histamine stimulation, compared to the parental HOS. Conversely, HM1OE cells tend to reduce mitochondrial Ca^2+^ uptake, after histamine induction (Fig Sup. 2b). Energetic profiling using Seahorse assays revealed a metabolic shift in HM1KO cells, characterized by an increased glycolytic activity and decreased oxidative phosphorylation (OXPHOS), consistent with the energetic consequences observed in mitoxantrone-treated HOS cells. In contrast, enhanced OXPHOS and reduced glycolysis were reported for HM1OE cells (Fig Sup. 2c). Transmission electron microscopy revealed that both HM1KO and HM1OE cell lines exhibited abnormal cristae structure and mitochondrial morphology (Fig Sup. 2d). Collectively, these results suggest that a decrease in MICU1 expression level directly modulates mitochondrial Ca^2+^ homeostasis, energetic metabolism, and mitochondrial ultrastructure. Therefore, the effects of mitoxantrone treatment observed in osteosarcoma cells might be partially mediated through MICU1.

### Metabolomic profiling of HOS cells following mitoxantrone treatment reveals metabolic vulnerability

To further dissect the compensatory metabolic reprogramming induced by mitoxantrone, evaluation of predefined metabolites associated with major metabolic pathways was conducted in HOS cells after 72h of mitoxantrone treatment at the IC_50_. Metabolomic profiling was performed using UHPLC–MS/MS and GC–MS/MS platforms with subsequent quantitative data processing and enrichment analysis via MetaboAnalyst 6.0. Analysis of differentially expressed metabolites suggested enhanced lipid metabolism, as evidenced by increased levels of 4 long-chain acylcarnitines (i.e. tetracosenoic acid, docosadienoic acid, myristoleic acid, docosatrienoic acid and docosenoic acid) (Fig. 4a). In line with these results, we next conducted the Seahorse Substrate Oxidation Stress test to assess the dependence of HOS cells on β-oxidation upon mitoxantrone treatment. In the HOS control cells, injection of the Carnitine palmitoyltransferase (CPT1) inhibitor - etomoxir led to a decrease in OCR value, suggesting a contribution of fatty acid oxidation (FAO) to mitochondrial respiration in this cell line. Upon 72h of mitoxantrone treatment, HOS cells displayed no decrease in OCR and were insensitive to etomoxir (Fig. 4b). These findings indicate that mitoxantrone has induced a metabolic reprogramming, in which FAO no longer significantly contributed to mitochondrial respiration, despite the accumulation of lipid intermediates.

**Figure 4.**
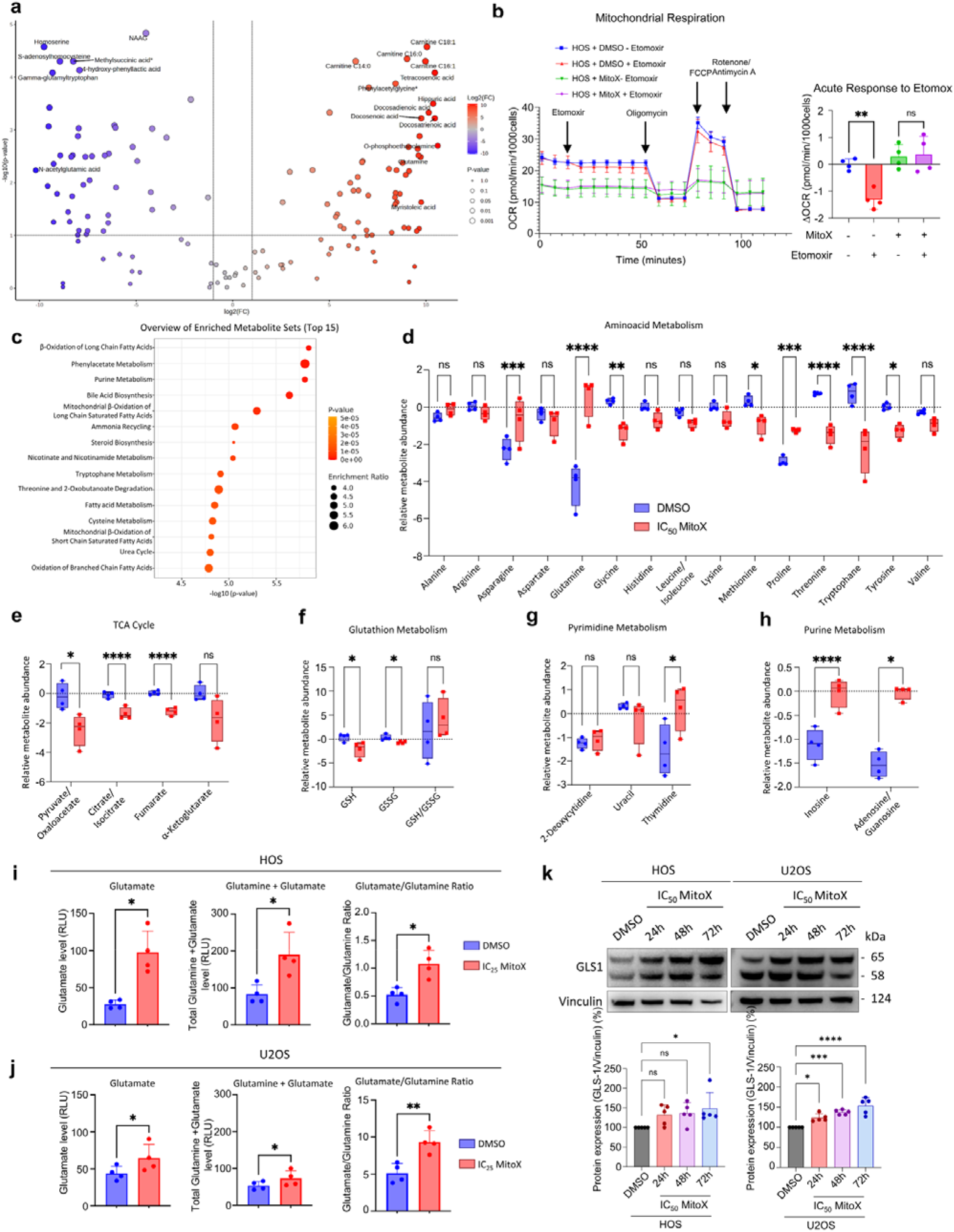
Metabolomic analysis of mitoxantrone-treated HOS cells shows a metabolic reprogramming toward glutamine accumulation and enhanced nucleotide synthesis. **a**, Volcano plot of metabolomic profiling data generated with MetaboAnalyst 6.0. Data were processed using the Statistical Analysis and Enrichment Analysis modules, with log_10_ transformation applied for normalization. The thresholds were set at –log_10_(p-value) = 1 for statistical significance and log_2_(fold change) = 1 for differential abundance. **b**, Seahorse XF Long Chain Fatty Acid Oxidation Stress Test graph displaying the oxygen consumption rate (OCR; pmol/min/1.000 cells) over time in HOS cells. The test was conducted using the following compounds: etomoxir (16µM), oligomycin (2.5 µM), FCCP (1 µM), and rotenone/antimycin A (0.5 µM). The acute response to etomoxir is shown as the difference in OCR measured immediately before and after etomoxir injection, reflecting the contribution of long-chain fatty acid oxidation to mitochondrial respiration. Data represent the mean ± SD from four independent experiments. Statistical significance was assessed using ANOVA with Sidak’s multiple comparison test correction, with significance levels indicated as follows: ns, not significant; p < 0.0332 (*); p < 0.0021 (**); p < 0.0002 (***); p < 0.0001 (****). **c**, Enrichment analysis of metabolites that differ significantly between 0.0005% DMSO as negative control and IC_50_ mitoxantrone-treated HOS cells following 72h of treatment. The enrichment ratio represents the observed number of metabolites in a specific metabolic pathway divided by the expected number. Metabolic pathways are ranked by p-value, with the most significant pathways at the top. **d** Relative abundance of amino acids showing significant alterations upon IC_50_ mitoxantrone treatment compared to 0.0005% DMSO control. Data represent relative metabolite abundance expressed as normalized log_2_-transformed values (areacorrLog2Cen). Data represent the mean ± SD from four independent experiments. Statistical significance was assessed using ANOVA with Sidak’s multiple comparison test correction, with significance levels indicated as follows: ns, not significant; p < 0.0332 (*); p < 0.0021 (**); p < 0.0002 (***);p < 0.0001 (****). **e-h**, Analysis of different metabolites involved in TCA cycle (pyruvate/oxaloacetate, citrate/isocitrate, fumarate, and α-ketoglutarate) (e); glutathione metabolism (GSH, GSSG and GSH/GSSG ratio) (f); pyrimidine metabolism (uracil, thymidine, and 2-deoxycytidine) (g) purine metabolism (adenosine/guanosine and inosine) (h). Data represent relative metabolite abundance expressed as normalized log_2_-transformed values (areacorrLog2Cen). Data represent the mean ± SD from four independent experiments. Statistical significance was assessed using ANOVA with Sidak’s multiple comparison test correction, with significance levels indicated as follows: ns, not significant; p < 0.0332 (*); p < 0.0021 (**); p < 0.0002 (***); p < 0.0001 (****). **i-j**, Mitoxantrone-induced increase in glutamate/glutamine ratio. Glutamate, total glutamate + glutamine, and glutamate/glutamine ratio of controls or mitoxantrone-treated HOS and U2OS cells. Cells were treated with IC_25_ of mitoxantrone or DMSO as a negative control for 72h, and levels of glutamate and total glutamate + glutamine were semi-quantified using the Glutamine/Glutamate-Glo™ Assay (Promega, #J8021). Glutamate/glutamine ratio was calculated accordingly. Data are presented as the mean ± SD from four independent experiments (n = 4). Statistical significance was assessed using an unpaired two-tailed Student’s t-test. Significance levels are indicated as follows: not significant (ns); p < 0.0332 (*); p < 0.0021 (**); p < 0.0002 (***); and p < 0.0001 (****). **k**, Upregulation of glutaminase 1 (GLS1) after mitoxantrone treatment. Western blot analysis showing the expression levels of GLS1 in response to either 0.0005% DMSO as a negative control or IC_50_ of mitoxantrone treatment over time (24, 48 and 72h). Vinculin was used as loading control. Data are presented as the mean ± SD from five independent experiments. Statistical significance was assessed using ANOVA with Sidak’s correction for multiple comparisons. Significance levels are indicated as follows: not significant (ns), p < 0.0332 (*), p < 0.0021 (**), p < 0.0002 (***), and p < 0.0001 (****).

In addition to changes in lipid metabolism, downregulation was observed in metabolites predominantly associated with glutamate-dependent processes, including N-acetyl-aspartyl-glutamate (NAAG), N-acetylglutamic acid, gamma-glutamyltryptophan, gamma-glutamyltyrosine, and S-adenosylhomocysteine. However, the volcano plot analysis revealed an increase in glutamine level, suggesting that this enriched glutamine was not directed to these glutamate-associated metabolites. (Fig. 4a). Enrichment analysis of differentially abundant metabolites revealed β-oxidation of fatty acids as one of the most enriched pathways (Fig. 4c), consistent with the observed increases in acylcarnitines and long-chain fatty acids. Purine metabolism was also enriched, suggesting a possible redirection of glutamine towards enhanced nucleotide biosynthesis upon mitoxantrone treatment. Moreover, comprehensive analysis showed decreased levels in the majority of measured amino acids, except for glutamine, proline, and asparagine, further highlighting the significance of glutamine in this metabolic reprogramming profile (Fig. 4d).

Cancer cells frequently exhibit glutamine addiction and utilize glutamine for various cellular activities such as energy production, fatty acid synthesis, regulation of redox homeostasis, and nucleotide synthesis ^27 28^. Therefore, we examined the metabolites associated with each of these pathways. First, glutamine-derived glutamate can be converted into *α*-ketoglutarate (*α*KG) that enters the tricarboxylic acid (TCA) cycle. This fueling of the TCA cycle either results in NADH and FADH_2_ production, which contributes directly to ATP synthesis via OXPHOS, or supports citrate production in fatty acid synthesis. However, in mitoxantrone-treated samples, decreased levels were observed in substrates of the TCA cycle, including pyruvate/oxaloacetate, citrate/isocitrate, fumarate, and *α*KG (Fig. 4e). Another role of glutamine-derived glutamate could be to modulate redox homeostasis via the synthesis of glutathione, a glutamate-cysteine-glycine tripeptide that neutralizes free radicals through redox reaction. Nevertheless, a reduction was reported for both reduced glutathione (GSH) and oxidized glutathione (GSSG) levels, concomitantly with an insignificant change in GSH/GSSG ratio, suggesting that glutamine was not re-directed to this process (Fig. 4f). From these results, it could be reasoned that glutamine was likely not entering the TCA cycle, nor contributing to glutathione biosynthesis. Lastly, as glutamine-derived glutamate serves also as a nitrogen donor and a source of carbon in nucleotide metabolism, we checked the intermediate metabolites of nucleotide synthesis. Here, significant upregulation was shown for thymidine, adenosine/guanosine, and inosine (Fig. 4g, h). Combined with the enriched nucleotide metabolism above, this indicates a potential redirection of glutamine to nucleotide metabolism in HOS cells after mitoxantrone treatment. Overall, these metabolomic alterations highlighted mitoxantrone-induced metabolic reprogramming signature, whereas the increase in glutamine level might contribute to nucleotide biosynthesis.

As a limit in our study, the glutamate quantification in our metabolomic profiling was not interpretable and did not allow an accurate estimation of glutamine to glutamate conversion. This led us to conduct a bio-luminescent assay to measure both glutamate and glutamine levels, as well as the glutamate/glutamine ratio in HOS (Fig. 4i) and U2OS (Fig. 4j) cells. Both substrates were increased in mitoxantrone-treated osteosarcoma cells compared to their controls. Additionally, the glutamate/ glutamine ratio approximately doubled in both cell lines, indicating an enhanced conversion of glutamine to glutamate (Fig. 4i, j). As glutamine is converted to glutamate by the enzyme glutaminase 1 (GLS1), we assessed GLS1 protein expression levels following mitoxantrone treatment at different time points (24, 48, and 72h). Increased GLS1 protein levels were found in both HOS and U2OS cell lines starting at 24h post-treatment (Fig. 4k). These results highlight the effects of mitoxantrone on glutamine-metabolism involving the GLS1 enzyme in osteosarcoma cell lines, without excluding the potential involvement of other factors in glutamine metabolism.

### Targeting metabolic vulnerability induced by mitoxantrone treatment - Synergism between telaglenastat and mitoxantrone

Based on the above results, we observed that glutamine accumulation and increased GLS1 activity were the metabolic consequences upon mitoxantrone treatment in osteosarcoma cells. This prompted us to evaluate the combined therapeutic effects of mitoxantrone with telaglenastat, a potent GLS1 inhibitor, which is currently under evaluation in clinical trials. Telaglenastat treatment of HOS cells resulted in moderate cytotoxicity with an IC_50_ of 13μM, while U2OS, Saos2 and 143B were insensitive to doses up to 100μM (Fig. 5a). To analyse the *in vitro* effects of mitoxantrone with telaglenastat, we used the Highest Single Agent (HSA) model in the SynergyFinder software. U2OS, HOS, and Saos2 cells showed synergistic interactions (i.e. scores > 10), with HSA scores of 19.22, 11.99, and 11.29, respectively, and 143B cells exhibited a score of 8.404, which was considered as additive (Fig. 5b).

**Figure 5.**
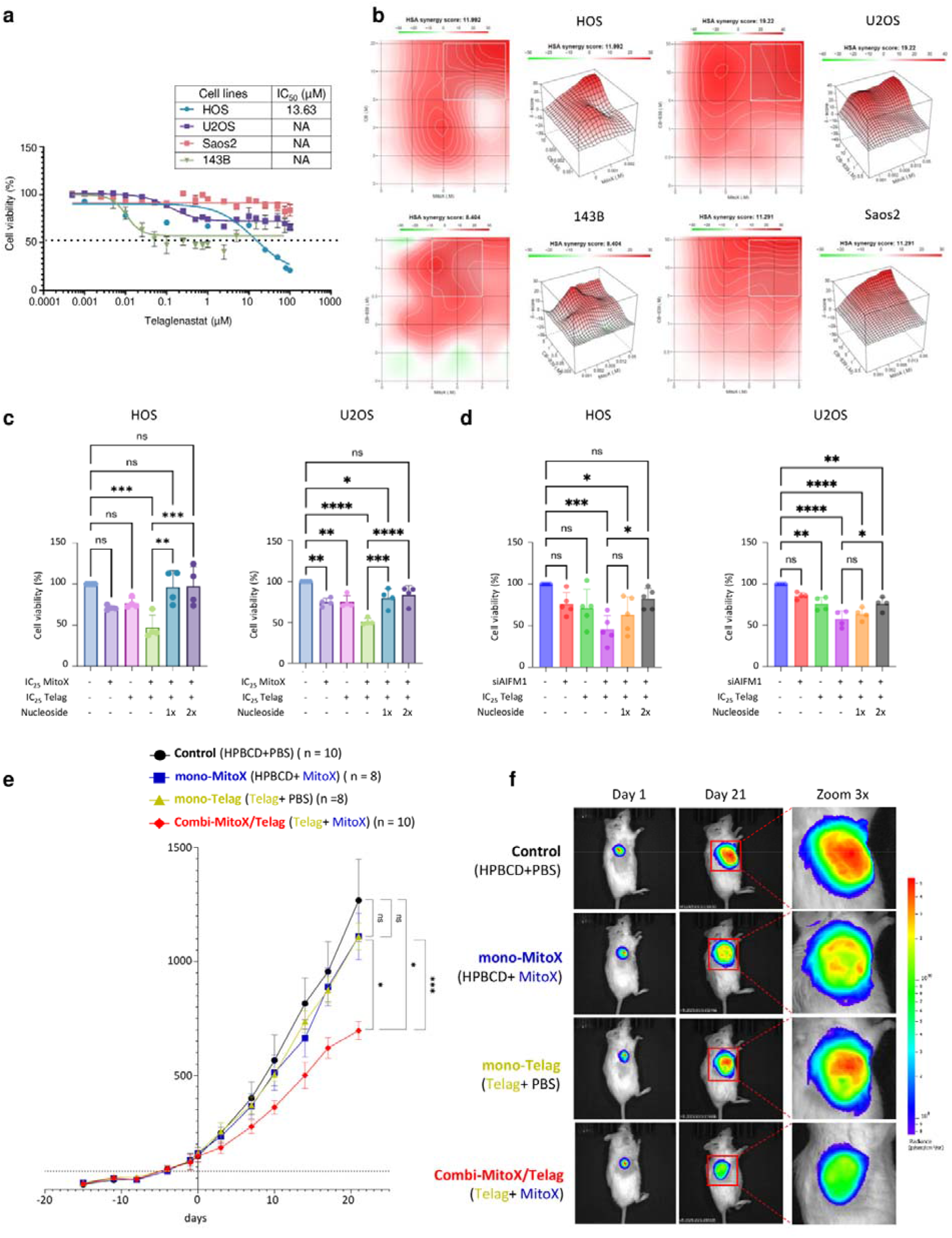
Targeting mitoxantrone-induced metabolic vulnerability through GLS1 inhibition: synergistic effects of telaglenastat in in *vitro* and *in vivo* osteosarcoma models. **a**, Cell viability of 4 pediatric osteosarcoma cell lines HOS, U2OS, SAOS2, 143B treated with different concentrations of telaglenastat, measured using an LDH assay. Dose-response curves represent cell viability at increasing concentrations of telaglenastat after 72h of treatment. 0.0005% DMSO was used as the solvent of mitoxantrone and as control for calculation of percentage of viability. Data were shown as mean ± SD from > 2 individual replicates. **b**, Illustrative 2D and 3D maps with overall HSA synergy score for mitoxantrone/telaglenastat combination in the osteosarcoma cell lines U2OS, HOS, SAOS2, 143B, obtained using SynergyFinder tool (https://synergyfinder.fimm.fi/). Cells were treated with varying concentrations of both drugs in a matrix form. Cell viability was measured using LDH assay and input onto the SynergyFinder platform. Red, white, and green areas correspond to synergism (score > 10), additivity (-10 < score < 10) and antagonism (score <-10), respectively. **c**, Effects of mitoxantrone/telaglenastat combination on nucleotide biosynthesis. Combination-treated (IC_25_ mitoxantrone, IC_25_ telaglenastat) cells were supplemented with nucleoside at 1x (30 μM each of cytidine, guanosine, uridine, adenosine and 10 μM thymidine) or 2x. Cell viability was measured using LDH assay. Data represent the mean±SD from four independent experiments. Statistical significance was assessed using ANOVA with Sidak’s multiple comparison test correction, with significance levels indicated as follows: ns, not significant; p < 0.0332 (*); p < 0.0021 (**); p < 0.0002 (***);p < 0.0001 (****). **d**, Involvement of AIF in mitoxantrone/telaglenastat synergism through selective AIF silencing and targeting nucleotide biosynthesis. HOS and U2OS cells were treated with siAIF for 48h prior to treatment with IC_25_ of telaglenastat and addition of nucleoside at 1x (30 μM each of cytidine, guanosine, uridine, adenosine and 10μM thymidine) or 2x. Data represent the mean±SD from at least four independent experiments. Statistical significance was assessed using ANOVA with Sidak’s multiple comparison test correction, with significance levels indicated as follows: ns, not significant; p < 0.0332 (*); p < 0.0021 (**); p < 0.0002 (***); p < 0.0001 (****). **e**, Tumor volume curves of U2OS-engrafted mice treated with PBS + HPBCD (Vehicle; n = 10) or mitoxantrone + HPBCD (mono-mitoxantrone; n = 8) or PBS + telaglenastat (mono-telaglenastat; n = 8) or mitoxantrone + telaglenastat (combi-mitoxantrone/telaglenastat; n = 10). Data were shown as mean ± standard error of mean (SEM) and statistical significance was determined by Two-way ANOVA (mixed model). Significance levels are indicated as follows: ns, not significant; p < 0.0332 (*); p < 0.0021 (**); p < 0.0002 (***); p < 0.0001 (****). **f**, Representative *in-vivo* bioluminescence images of U2OS-Luc/mkate2-mice treated with PBS + HPBCD (Vehicle), mitoxantrone + HPBCD (mono-mitoxantrone), PBS + Telaglenastat (mono-telaglenastat), or mitoxantrone + telaglenastat (combi-mitoxantrone/telaglenastat).

To explore the importance of glutamine accumulation in nucleotide synthesis in cell response to mitoxantrone treatment and the strategy to combine it with telaglenastat, we performed a nucleoside rescue assay. The addition of the nucleosides cytidine, guanosine, uridine, adenosine, thymidine to mitoxantrone/telaglenastat-treated HOS and U2OS cells reversed the cytotoxic effects observed in unsupplemented conditions. Furthermore, siRNA-mediated knockdown of AIF resulted in reduced cell viability when combined with telaglenastat, thereby mimicking its inhibitory effect. Moreover, nucleoside supplementation was able to restore cell growth (Fig. 5d), supporting that the cytotoxic effect of mitoxantrone was mediated via selective inhibition of the AIF/CHCHD4 complex and its involvement in this adaptive mechanism (Fig. 5d).

Finally, to evaluate the anti-tumor effects of both agents alone and their combination *in vivo*, subcutaneous U2OS-Luc/mkate2-engrafted NSG mice were treated at advanced tumor stage four weeks after tumor cell engraftment. Mice bearing tumors of 80-120 mm^3^ were randomly assigned into four treatment groups: vehicle (n = 10), mitoxantrone 1mg/kg day 1 and 4 (n = 8), telaglenastat 200mg/kg b.i.d day 1 to day 21 (n = 8), and mitoxantrone/telaglenastat combination in equivalent doses and schedules (n = 10). Treatment continued for three weeks, during which tumor growth and body weight were monitored. At the end of the treatment, we observed that neither mitoxantrone nor telaglenastat monotherapy at these low doses significantly reduced tumor growth compared with vehicle controls, after 3 weeks of treatment. In contrast, the combination of mitoxantrone/telaglenastat (696.6 ± 39.45 mm^3^, mean ± SEM) significantly inhibited tumor progression, with tumors displaying significantly smaller volumes, compared to either the vehicle group (1268 ± 180.6 mm^3^, mean ± SEM) or mitoxantrone monotherapy (1110 ± 101.9 mm^3^, mean ± SEM) or telaglenastat monotherapy group (1110 ± 58.57 mm^3^, mean ± SEM) (Fig. 5e, f). These findings reinforced the synergistic interaction previously observed *in vitro*, demonstrating that mitoxantrone and telaglenastat combination effectively suppressed osteosarcoma growth *in vivo* at doses that were individually insensitive.

## Discussion

Here, we show a new mechanism of action of mitoxantrone, a chemotherapeutic agent, that induces a metabolic vulnerability that proved to be instrumental for combination with teleglanastat, a GLS1 inhibitor. This highlights the pharmacological potential of interrupting the AIF/CHCHD4 complex in OS.

Combining *in vitro* high-throughput screening and *in silico* analysis, we revealed mitoxantrone as an inhibitor of AIF/CHCHD4 interaction, potentially through competitive binding to the NAD-binding site of AIF. As publicly available data showed that high expression of *AIFM1* and *CHCHD4* mRNA correlated with poor overall survival in pediatric OS and Ewing sarcoma, while osteosarcoma patients in two trials MAPPYACTS and OS2006 exhibited significant levels of *AIF* and *CHCHD4* ^*8*^, we decided to explore the impact of mitoxantrone-induced inhibition of AIF/CHCHD4 interaction in osteosarcoma models. Treatment with mitoxantrone resulted in cytotoxicity of osteosarcoma cell lines, including resistant ones, and of PDX cell cultures derived from refractory and relapsed osteosarcoma patients. The cell death was preceded by profound changes in mitochondrial structure, bioenergetics and Ca^2+^ accumulation. Mitoxantrone also induced DNA damage, as evidenced by an increase in phosphorylated histone H2AX - a marker of double-strand breaks, thus confirming that its classical mechanism of action was actionable in our osteosarcoma models (Fig Supp. 1d). Therefore, our findings expand our knowledge of the anticancer properties of mitoxantrone beyond DNA intercalation and inhibition of topoisomerase II ^10 29^.

Mitoxantrone is currently approved for the treatment of patients with leukemias, advanced breast cancer, lymphomas and prostate cancer ^10 29 30 31 32^. Here, mitoxantrone exhibited high efficacy in secondary cultured PDX *in vitro* models collected from MAPPYACTS cohort of relapsed, resistant patients, compared to currently used conventional chemotherapies. This novel use of mitoxantrone suggests that disrupting the AIF/CHCHD4 interaction could be a potential therapeutic strategy for treatment of chemoresistant osteosarcoma, which would expand its clinical utility. Nevertheless, further studies are required to validate and better characterize this approach.Mechanistically, we demonstrated that mitoxantrone specifically disrupted the AIF/CHCHD4 complex, as shown by co-immunoprecipitation and AIF knockdown. This disruption resulted in decreased levels of several downstream mitochondrial substrates including mETC subunits CI, CIV, and MICU1, thereby contributing to associated impairment of OXPHOS and altered mitochondrial Ca^2+^ uptake, respectively. Of note, previous studies have reported that mitoxantrone reduces mitochondrial Ca^2+^ uptake by inhibiting MCU. On the other hand, our observations suggest an alternative contribution from MICU1, which may reflect cell line–dependent differences ^33 34^. In an attempt to understand the link between mitoxantrone-induced decrease in MICU1 and metabolic consequences in osteosarcoma, we have generated MICU1-deficient HOS cells and reported similar phenotypes when compared to mitoxantrone-treated HOS cells. Moreover, the glutamine accumulation was also observed in metabolomic data analysis while comparing HM1KO cells to parental HOS cells (Fig Supp. 2e). These findings suggest that, in our model, mitoxantrone might act directly through MICU1, without excluding any effect on MCU or other factors, to alter mitochondrial Ca^2+^ accumulation and mediate some of the metabolic perturbations induced by mitoxantrone.

Interestingly, mitoxantrone-induced reduction in OCR and proton leak were accompanied by ΔΨm hyperpolarization, indicating a high-potential, low-respiration state. Several scenarios could explain this observation: i) reduced forward mode of complex V; ii) activation of reverse mode of complex V; iii) decreased availability of ADP in the mitochondrial matrix ^35 36 37^. Therefore, in the future, different specific inhibitors could be used to validate those hypotheses such as BTB06584 (reverse-mode ATP synthase inhibitor) ^38^ and carboxyatractyloside (adenine nucleotide translocase (ANT) inhibitor) ^39^ Consistent with our findings, it has been demonstrated that 24h treatment with rotenone, an inhibitor of CI, leads to ΔΨm hyperpolarization in HEK293 cells ^40^. In this study, the authors argued that despite chronic CI inhibition, the mitochondrial matrix did not acidify, while CV exhibited reduced activity and was not triggered into reverse mode. Altogether, rotenone treatment created proton-based ΔΨm hyperpolarization, and mitoxantrone treatment might induce a similar phenomenon through CI downregulation^40^.

A novel concept in anti-cancer therapies is to target metabolic vulnerability ^41^. Our studies in the metabolomic profile of mitoxantrone-treated osteosarcoma cells revealed the induction of glutamine accumulation that potentially fuels nucleotide biosynthesis as a compensatory survival mechanism. Enhanced conversion of glutamine to glutamate and increased GLS1 protein expression level suggested a potential synergistic activity when combining mitoxantrone with GLS1 inhibitor. Among currently available inhibitors of glutamine metabolism, the GLS1 inhibitor telaglenastat has shown promising efficacy in preclinical studies and is currently in early clinical development as single agent and in combination therapies. Therefore, we selected it for the combination experiments with mitoxantrone and showed better tumor growth inhibition in xenograft mice models, compared to single agent treatment Our findings are in alignment with other studies showing mETC defects suppressing *de novo* purine synthesis and increasing dependence on purine salvage pathways ^42^. Since the AIF/CHCHD4 complex is required for mETC biogenesis, it is possible that mitoxantrone monotherapy indirectly diminishes nucleotide biosynthesis through downregulation of mETC complexes. Moreover, models of ovarian cancer treated with telaglenastat alone also exhibited perturbations in nucleotide metabolism ^43^. Therefore, the combination of telaglenastat with mitoxantrone in osteosarcoma treatment potentially amplifies this inhibitory effect on nucleotide biosynthesis. Altogether, these findings emphasize that targeting mitochondrial mETC alters energy production and induces biosynthetic reprogramming.

In therapy-induced senescent melanoma cells, CDK4/6 inhibition with palbociclib leads to GLS1 upregulation, creating a glutamine-addicted state that can be selectively inhibited with telaglenastat ^44^. Similarly, MAPK pathway inhibition in melanoma or *KRAS*-driven non-small-cell lung cancer (NSCLC) increases glutamine dependency, and the combination of selumetinib with telaglenastat shows robust synergy and tumor control ^45^. Resistance to BRAF inhibitors in melanoma has also been reported to drive glutamine dependence and mitochondrial reliance, providing a rationale for co-targeting GLS1 alongside MAPK signaling ^46^. These examples highlight that diverse perturbations can remodel tumor metabolism toward glutamine and nucleotide dependency, which can then be exploited by GLS1 inhibition. Thus, our findings position AIF/CHCHD4 inhibition by mitoxantrone as a novel mitochondrial target that induces osteosarcoma cells into a glutamine-nucleotide coping mechanism. More importantly, the similar scenario in other cancer contexts suggests that the metabolic vulnerability concept is not confined to osteosarcoma but represents a feasible therapeutic strategy across diverse cancer types. Collectively, our study has established a mechanistic link between AIF/CHCHD4 disruption and glutamine metabolic reprogramming and provided preclinical evidence that targeting AIF/CHCHD4 complex by mitoxantrone combined with GLS1 inhibitor telaglenastat could represent a therapeutic opportunity.

One of the limitations of our study is that the precise biophysical mechanism by which mitoxantrone binds to AIF remains to be elucidated by crystallography and/or cryo-EM. Second, the precise steps in which glutamine contributes to nucleotide synthesis upon mitoxantrone treatment are not yet defined. Third, while our *in vivo* results showed a partial inhibition of tumor growth, they were limited to one osteosarcoma xenograft model, and further validation in additional PDX and orthotopic systems is warranted.

In conclusion, the identification of mitoxantrone as an inhibitor of the AIF/CHCHD4 complex not only provides a new insight into its mechanisms of action but also opens new avenues for the development of next-generation small molecules specifically targeting this mitochondrial complex and for rational combinatory strategies to overcome resistance in cancer treatment.

## Acknowledgments

CB research was supported by grants from the French National Cancer Institute “INCa 2017-1-PL BIO-08” and “2021 - 167/ INCA_16344” and the Société française de lutte contre les cancers et les leucémies de l’enfant et de l’adolescent (SFCE), grant number ECS 20, PHC PESSOA, N°49163TL and the Groupement des Entreprises Françaises dans la lutte contre le Cancer [2023-2024]. BG work including the MAPPYACTS trial was supported by grants from INCa through the PHRC “INCa-DGOS_8519” MERRI, Fondation ARC, Association Imagine for Margo, Fédération Enfants et Santé, SFCE, and Dell. The MAPPYACTS PDX establishment was supported by grants from Fondation Gustave Roussy; Fédération Enfants Cancers et Santé, Société Française de lutte contre les Cancers et les leucémies de l’Enfant et l’adolescent (SFCE), Association AREMIG and Thibault BRIET; Parrainage médecin-chercheur of Gustave Roussy. AM and BG were supported by the “Parrainage médecin-chercheur” of Gustave Roussy. The work has been produced in collaboration with the BoOST-dataS consortium, coordinated by Gustave Roussy and including the Toulouse University Hospital, the UMR1238 unit of the University of Nantes, the Léon Bérard centre, the Curie Institute, Unicancer, the Strasbourg University Hospital & University, the CRBM of the University of Montpellier and the Oscar Lambret centre, and with the support of the French National Cancer Institute. Sylvie Souquère (EM platform) from UMS AMMICA, Gustave Roussy, Villejuif, Manon Cruette (INSERM U1180, Univ Paris-Saclay, Châtenay-Malabry), Delphine Courilleau (CIBLOT platform, Univ Paris-Saclay, Gif Sur Yvette), Magali Noiray (I2BC, Univ Paris-Saclay, Gif sur Yvette), Dawei Liu (UMR9018, Gustave Roussy) are acknowledged for their technical help. Dr Jan Koster for R2: Genomics Analysis and Visualization platform. Figures were created with BioRender.com

## Conflicts of interest

GK has been holding research contracts with Daiichi Sankyo, Eleor, Kaleido, Lytix Pharma, PharmaMar, Osasuna Therapeutics, Samsara Therapeutics, Sanofi, Sutro, Tollys, and Vascage. GK is on the Board of Directors of the Bristol Myers Squibb Foundation France. GK is a scientific co-founder of everImmune, Osasuna Therapeutics, Samsara Therapeutics and Therafast Bio. GK is in the scientific advisory boards of Centenara Labs (formerly Rejuveron Life Sciences), Hevolution, and Institut Servier. GK is the inventor of patents covering therapeutic targeting of aging, cancer, cystic fibrosis and metabolic disorders. Among these, patents were licensed to Bayer (WO2014020041-A1, WO2014020043-A1), Bristoll-Myers Squibb (WO2008057863-A1), Osasuna Therapeutics (WO2019057742A1), PharmaMar (WO2022049270A1 and WO2022048775-A1), Raptor Pharmaceuticals (EP2664326-A1), Samsara Therapeutics (GB202017553D0), and Therafast Bio (EP3684471A1). GK’wife, Laurence Zitvogel, has held research contracts with Glaxo Smyth Kline, Incyte, Lytix, Kaleido, Innovate Pharma, Daiichi Sankyo, Pilege, Merus, Transgene, 9 m, Tusk and Roche, was on the on the Board of Directors of Transgene, is a cofounder of everImmune, and holds patents covering the treatment of cancer and the therapeutic manipulation of the microbiota. Among these, patents were licensed to everImmune (US20210346438A1, US20200360449A1) and Transgene (US20200376052A1). GK’s brother, Romano Kroemer, was an employee of Sanofi and now consults for Boehringer-Ingelheim. The funders had no role in the design of the study; in the writing of the manuscript, or in the decision to publish the results.

Nathalie Gaspar has got consultant activity for EISAI and IPSEN, and participate to advisory board for ABBVIE, MERCKS, Y-MABS, Daiichi Sankyo/AstraZeneca.

## Notes

### Competing Interest Statement

The authors have declared no competing interest.

## References

1. Gaspar, N. et al. Recent advances in understanding osteosarcoma and emerging therapies. Faculty Reviews 9, 18 (2020).

2. Marchandet, L. et al. Mechanisms of Resistance to Conventional Therapies for Osteosarcoma. Cancers 13, 683 (2021).

3. Giang, A.-H. et al. Mitochondrial dysfunction and permeability transition in osteosarcoma cells showing the Warburg effect. J Biol Chem 288, 33303–33311 (2013).

4. Palorini, R. et al. Energy Metabolism Characterization of a Novel Cancer Stem Cell-Like Line 3AB-OS. Journal of Cellular Biochemistry 115, 368–379 (2014).

5. Ren, L. et al. Glutaminase-1 (GLS1) inhibition limits metastatic progression in osteosarcoma. Cancer Metab 8, 4 (2020).

6. Porporato, P. E., Filigheddu, N., Pedro, J. M. B.-S., Kroemer, G. & Galluzzi, L. Mitochondrial metabolism and cancer. Cell Res 28, 265–280 (2018).

7. Chen, F. et al. The role of mitochondria in tumor metastasis and advances in mitochondria-targeted cancer therapy. Cancer Metastasis Rev 43, 1419–1443 (2024).

8. Lai, H. T. et al. Engineered flavonoid disrupts mitochondrial AIF/CHCHD4 complex for targeted cancer therapy. 2025.03.13.642976 Preprint at 10.1101/2025.03.13.642976 (2025).

9. Reinhardt, C. et al. AIF meets the CHCHD4/Mia40-dependent mitochondrial import pathway. Biochim Biophys Acta Mol Basis Dis 1866, 165746 (2020).

10. Evison, B. J., Sleebs, B. E., Watson, K. G., Phillips, D. R. & Cutts, S. M. Mitoxantrone, More than Just Another Topoisomerase II Poison. Medicinal Research Reviews 36, 248–299 (2016).

11. Hangen, E. et al. Interaction between AIF and CHCHD4 Regulates Respiratory Chain Biogenesis. Mol Cell 58, 1001–1014 (2015).

12. Ye, H. et al. DNA binding is required for the apoptogenic action of apoptosis inducing factor. Nat Struct Biol 9, 680–684 (2002).

13. Waterhouse, A. et al. SWISS-MODEL: homology modelling of protein structures and complexes. Nucleic Acids Res 46, W296–W303 (2018).

14. Abraham, M. J. et al. GROMACS: High performance molecular simulations through multi-level parallelism from laptops to supercomputers. SoftwareX 1–2, 19–25 (2015).

15. Lindorff-Larsen, K. et al. Improved side-chain torsion potentials for the Amber ff99SB protein force field. Proteins 78, 1950–1958 (2010).

16. Jorgensen, W. L., Chandrasekhar, J., Madura, J. D., Impey, R. W. & Klein, M. L. Comparison of simple potential functions for simulating liquid water. The Journal of Chemical Physics 79, 926–935 (1983).

17. Trott, O. & Olson, A. J. AutoDock Vina: improving the speed and accuracy of docking with a new scoring function, efficient optimization, and multithreading. J Comput Chem 31, 455–461 (2010).

18. Missiaglia, E. et al. PAX3/FOXO1 fusion gene status is the key prognostic molecular marker in rhabdomyosarcoma and significantly improves current risk stratification. J Clin Oncol 30, 1670–1677 (2012).

19. Marques da Costa, M. E. et al. In-Vitro and In-Vivo Establishment and Characterization of Bioluminescent Orthotopic Chemotherapy-Resistant Human Osteosarcoma Models in NSG Mice. Cancers 11, 997 (2019).

20. da Costa, M. E. M. et al. Longitudinal characterization of primary osteosarcoma and derived subcutaneous and orthotopic relapsed patient-derived xenograft models. Front Oncol 13, 1166063 (2023).

21. Canoy, R. J. et al. Specificity of cancer-related chromosomal translocations is linked to proximity after the DNA double-strand break and subsequent selection. NAR Cancer 5, zcad049 (2023).

22. Germini, D. et al. A One-Step PCR-Based Assay to Evaluate the Efficiency and Precision of Genomic DNA-Editing Tools. Mol Ther Methods Clin Dev 5, 43–50 (2017).

23. Berlanga, P. et al. The European MAPPYACTS Trial: Precision Medicine Program in Pediatric and Adolescent Patients with Recurrent Malignancies. Cancer Discovery OF1–OF16 (2022) doi:10.1158/2159-8290.CD-21-1136.

24. Marchais, A. et al. Immune Infiltrate and Tumor Microenvironment Transcriptional Programs Stratify Pediatric Osteosarcoma into Prognostic Groups at Diagnosis. Cancer Res 82, 974–985 (2022).

25. Skinner, K. T., Palkar, A. M. & Hong, A. L. Genetics of ABCB1 in Cancer. Cancers (Basel) 15, 4236 (2023).

26. Kamath, N. et al. Overexpression of P-glycoprotein and alterations in topoisomerase II in P388 mouse leukemia cells selected in vivo for resistance to mitoxantrone. Biochem Pharmacol 44, 937–945 (1992).

27. Yang, L., Venneti, S. & Nagrath, D. Glutaminolysis: A Hallmark of Cancer Metabolism. Annu Rev Biomed Eng 19, 163–194 (2017).

28. Jin, J., Byun, J.-K.Choi, Y.-K. & Park, K.-G. Targeting glutamine metabolism as a therapeutic strategy for cancer. Exp Mol Med 55, 706–715 (2023).

29. Faulds, D., Balfour, J. A., Chrisp, P. & Langtry, H. D. Mitoxantrone. A review of its pharmacodynamic and pharmacokinetic properties, and therapeutic potential in the chemotherapy of cancer. Drugs 41, 400–449 (1991).

30. Shenkenberg, T. D. & Von Hoff, D. D. Mitoxantrone: a new anticancer drug with significant clinical activity. Ann Intern Med 105, 67–81 (1986).

31. Allegra, J. C. et al. A randomized trial comparing mitoxantrone with doxorubicin in patients with stage IV breast cancer. Invest New Drugs 3, 153–161 (1985).

32. Gherlinzoni, F. et al. Phase III comparative trial (m-BACOD v m-BNCOD) in the treatment of stage II to IV non-Hodgkin’s lymphomas with intermediate-or high-grade histology. Semin Oncol 17, 3–8; discussion 8-9 (1990).

33. Arduino, D. M. et al. Systematic Identification of MCU Modulators by Orthogonal Interspecies Chemical Screening. Mol Cell 67, 711–723.e7 (2017).

34. Zhang, Z. et al. Ruthenium 360 and mitoxantrone inhibit mitochondrial calcium uniporter channel to prevent liver steatosis induced by high-fat diet. Br J Pharmacol 179, 2678–2696 (2022).

35. Rego, A. C., Vesce, S. & Nicholls, D. G. The mechanism of mitochondrial membrane potential retention following release of cytochrome c in apoptotic GT1-7 neural cells. Cell Death Differ 8, 995–1003 (2001).

36. Acin-Perez, R. et al. Inhibition of ATP synthase reverse activity restores energy homeostasis in mitochondrial pathologies. The EMBO Journal 42, e111699 (2023).

37. Halestrap, A. P. & Brenner, C. The adenine nucleotide translocase: a central component of the mitochondrial permeability transition pore and key player in cell death. Curr Med Chem 10, 1507–1525 (2003).

38. Ivanes, F. et al. The compound BTB06584 is an IF1-dependent selective inhibitor of the mitochondrial F1Fo-ATPase. British Journal of Pharmacology 171, 4193–4206 (2014).

39. Shabalina, I. G., Kramarova, T. V., Nedergaard, J. & Cannon, B. Carboxyatractyloside effects on brown-fat mitochondria imply that the adenine nucleotide translocator isoforms ANT1 and ANT2 may be responsible for basal and fatty-acid-induced uncoupling respectively. Biochem J 399, 405–414 (2006).

40. Forkink, M. et al. Mitochondrial hyperpolarization during chronic complex I inhibition is sustained by low activity of complex II, III, IV and V. Biochimica et Biophysica Acta (BBA)-Bioenergetics 1837, 1247–1256 (2014).

41. Stine, Z. E., Schug, Z. T., Salvino, J. M. & Dang, C. V. Targeting cancer metabolism in the era of precision oncology. Nat Rev Drug Discov 21, 141–162 (2022).

42. Wu, Z. et al. Electron transport chain inhibition increases cellular dependence on purine transport and salvage. Cell Metab 36, 1504–1520.e9 (2024).

43. Shen, Y.-A. et al. Inhibition of the MYC-Regulated Glutaminase Metabolic Axis Is an Effective Synthetic Lethal Approach for Treating Chemoresistant Ovarian Cancers. Cancer Res 80, 4514–4526 (2020).

44. Kim, J. et al. Inhibition of glutaminase elicits senolysis in therapy-induced senescent melanoma cells. Cell Death Dis 15, 902 (2024).

45. Xia, M., Li, X., Diao, Y., Du, B. & Li, Y. Targeted inhibition of glutamine metabolism enhances the antitumor effect of selumetinib in KRAS-mutant NSCLC. Translational Oncology 14, 100920 (2021).

46. Baenke, F. et al. Resistance to BRAF inhibitors induces glutamine dependency in melanoma cells. Mol Oncol 10, 73–84 (2016).

