## Supplemental Data for "Combined inhibition of AIF/CHCHD4 interaction and GLS1 to exploit metabolic vulnerabilities in pediatric osteosarcoma"

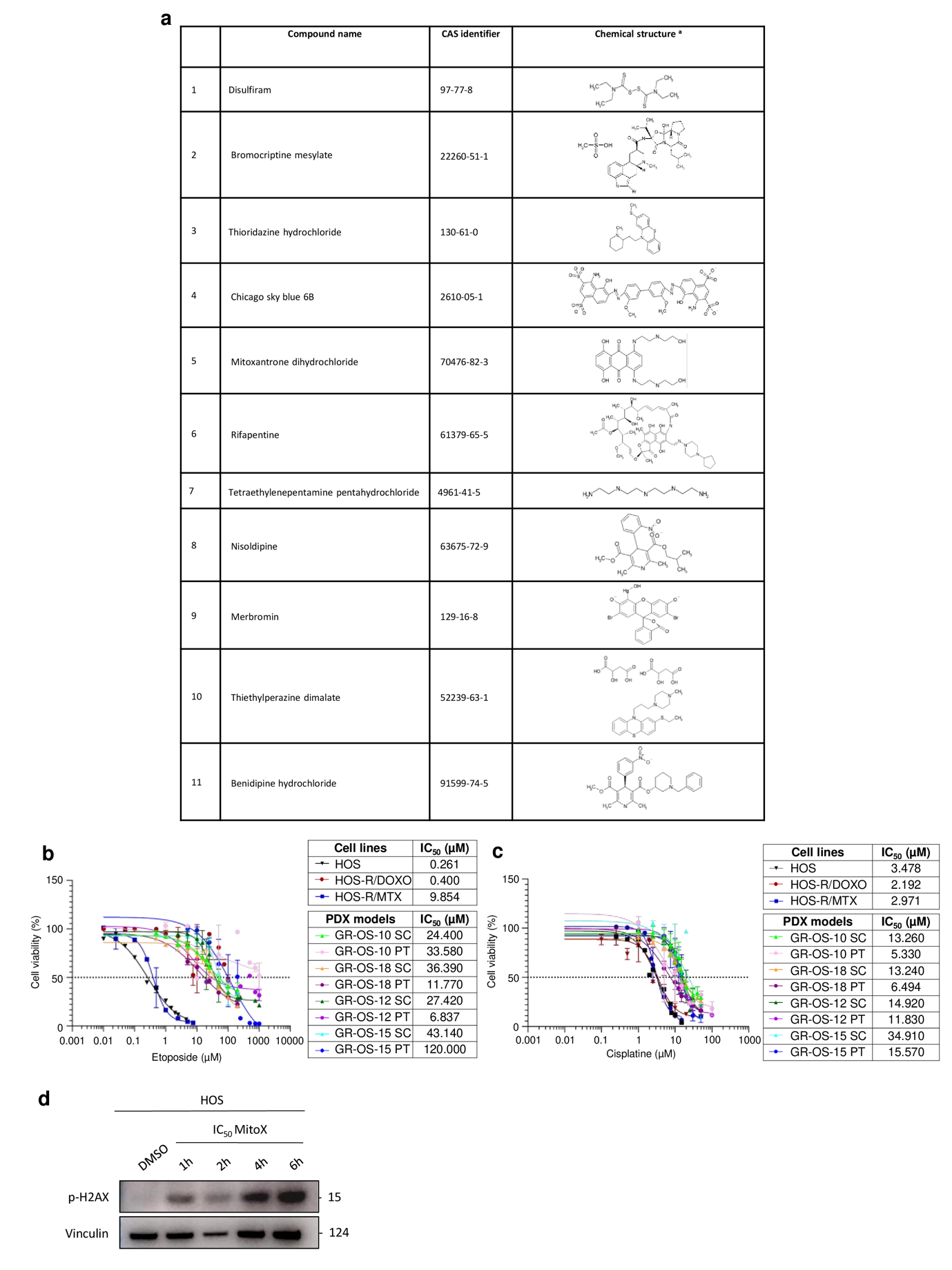

**Supplementary Figure 1.** **Selected compounds from Prestwick and DNA damage caused by mitoxantrone**

**a,** 11 compounds selected from Prestwick library by HTS Alpha screen, CAS identifiers and their chemical structure

**b,c,** Cell viability of 3 osteosarcoma cell lines and 8 secondary PDX cultures in vitro treated with different concentrations of etoposide (b) and cisplatin (c), and measured using an LDH assay. Dose-response curves represent cell viability at increasing concentrations of mitoxantrone after 72h of treatment and the IC₅₀ values were determined using GraphPad Prism software. Individual replicates are shown (n = 2). Data were shown as mean ± SD.

**d,** Time-dependent induction of DNA damage marker phospho-H2AX in HOS cells following mitoxantrone treatment

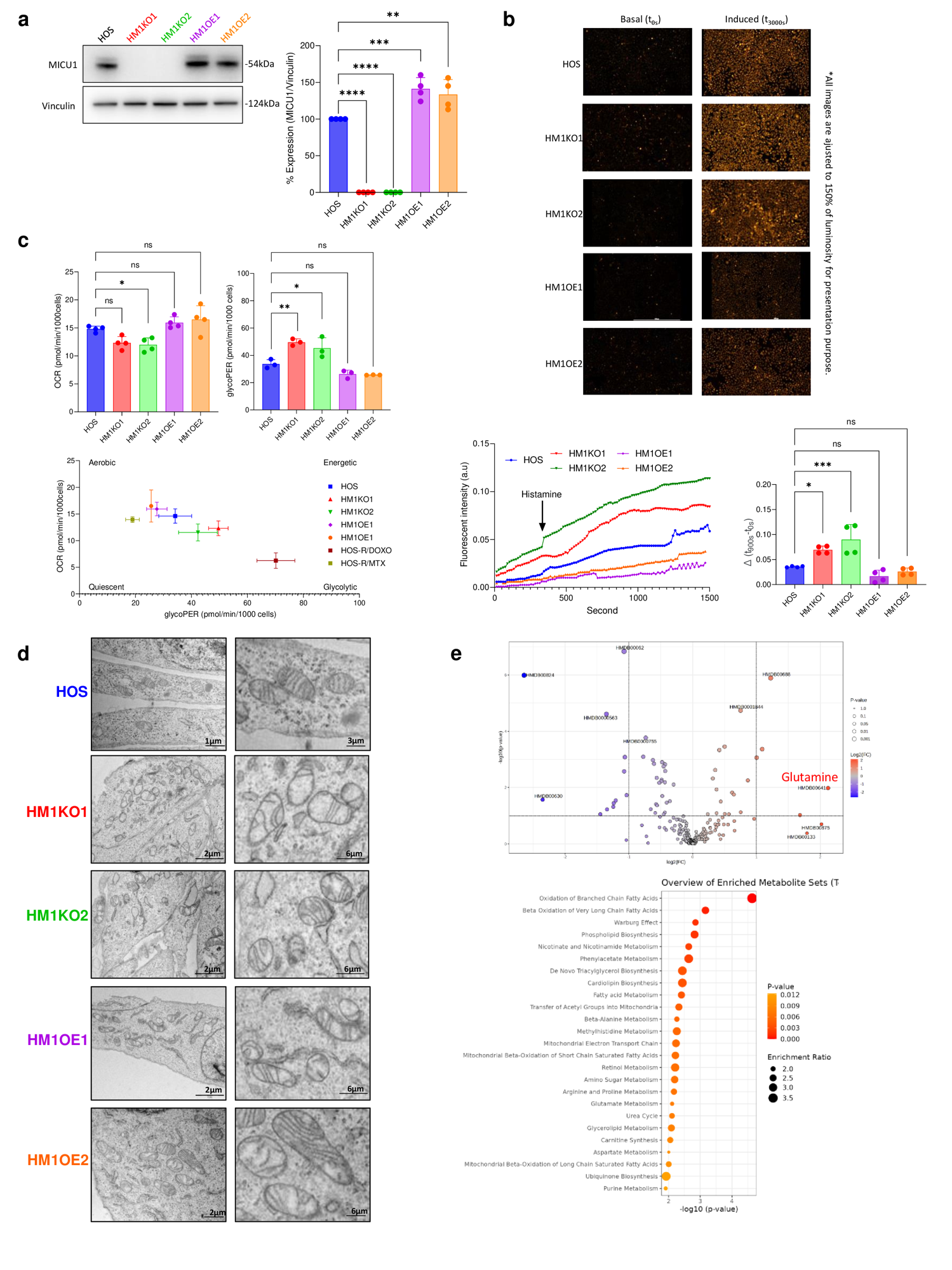

**Supplementary Figure 2. MICU1 modulation alters mitochondrial Ca²⁺ handling, ultrastructure, bioenergetics, and metabolic profiles in HOS cells.**

**a,** MICU1 protein expression in parental HOS cells and *MICU1* knockout (HM1KO1, HM1KO2) or overexpressing clones (HM1OE1, HM1OE2). Vinculin served as a loading control. Data are shown as mean ± SD; one-way ANOVA with Sidak’s multiple-comparison test (**p < 0.01; ****p < 0.0001; ns, not significant).

**b,** Histamine-induced mitochondrial Ca²⁺ uptake in different MICU1-modified OS cells measured using the fluorescent probe Rhod-2 AM. Cells were loaded with Rhod-2 AM (4 µM, 30 min) and Hoechst 33342 for nuclear counterstaining, and imaged using a Cytation 1 reader (Agilent). Baseline fluorescence was recorded for 5 min (interval = 2 s) before stimulation with histamine (100 µM) in the presence of extracellular CaCl₂ (200 mM), and changes in Rhod-2 fluorescence were monitored for an additional 30 min. Mean fluorescence intensity was normalized to Hoechst-positive nuclei.

**c,** Basal OCR respiration (pmol/min/1.000 cells) and basal glycolytic proton efflux rate (glycoPER) n different MICU1-modified OS cells. OCR and glycoPER measurements were normalized to cell counts determined by nuclei DAPI staining. Data represent the mean ± SD of at least three independent experiments. Statistical analysis was performed using ANOVA with Sidak’s correction for multiple comparisons, with significance levels indicated as follows: ns, not significant; p < 0.0332 (*); p < 0.0021 (**); p < 0.0002 (***);p < 0.0001 (****). Energy map was constructed using basal OCR and glycoPER values showing the balance between mitochondrial respiration (OCR) and glycolysis (glycoPER).

**d,** Transmission electron microscopy images of mitochondrial ultrastructure in HOS, MICU1 knockout (HM1KO1, HM1KO2), and MICU1 overexpressing (HM1OE1, HM1OE2) cells. Scale bars as indicated.

**e,** Metabolomic profiling of HM1KO1 cells. Volcano plot was processed using the Statistical Analysis and Enrichment Analysis modules, with log₁₀ transformation applied for normalization. The thresholds were set at –log₁₀(p-value) = 1 for statistical significance and log₂(fold change) = 1 for differential abundance. Enrichment analysis of metabolites that differ significantly between HOS and HM1KO1 cells. The enrichment ratio represents the observed number of metabolites in a specific metabolic pathway divided by the expected number. Metabolic pathways are ranked by p-value, with the most significant pathways at the top (glutamine indicated in red). Analysis was conducted using MetaboAnalyst 6.0.
